# Predicting Immunotherapy Response Through Chemotherapy-Induced Tumor Microenvironment Remodeling

**DOI:** 10.64898/2026.06.08.730822

**Authors:** Almog Angel, Zhongyang Lin, Michal Brunwesser, Dvir Aran

## Abstract

**Background:** Chemotherapy before immunotherapy improves outcomes in multiple cancers, but the benefit is heterogeneous: only some tumors undergo the immune remodeling that makes subsequent immunotherapy effective, and no biomarker identifies these patients prospectively, leading to uniform use of chemo-immunotherapy and avoidable toxicity.

**Methods:** A random forest model of tumor-microenvironment (TME) favorability, trained on 1,936 immunotherapy-treated patients, was applied to 334 paired pre-/post-chemotherapy biopsies across 19 studies to identify TME “converters,” from which a 50-gene baseline signature was derived. The signature was tested in IMvigor210 (the only cohort containing both treatment arms) and in five independent chemo-immunotherapy validation cohorts, and for specificity against 22 immunotherapy-only cohorts.

**Results:** Of 209 paired-biopsy patients with unfavorable baseline TME, 61 (29%) converted to a favorable state after chemotherapy; converters were indistinguishable from non-converters at baseline by cell composition, motivating a gene-level signature. In IMvigor210, the signature discriminated responders among chemotherapy-primed patients with unfavorable baseline TME (AUC = 0.80, 95% CI: 0.67–0.94) but showed no signal in the trial’s immunotherapy-only arm (interaction OR = 4.42, *p* = 0.006). Across five independent chemo-immunotherapy cohorts, responders consistently scored higher than non-responders, yielding a significant pooled effect by meta-analysis (pooled *g* = 0.59, 95% CI: 0.11–1.08, *p* = 0.017) and individual significance in two (LUD2015-005, *p* = 0.009; GSE165252, *p* = 0.02). A meta-analysis of 22 immunotherapy-only cohorts constrained any effect there to a small magnitude (Hedges’ *g* = 0.08, 95% CI: −0.08 to 0.23), and five established immune signatures did not reproduce this treatment-context specificity. Converter tumors harbored coordinated baseline programs of proliferative stress, innate and adaptive immune readiness, and functional vasculature.

**Conclusions:** A baseline transcriptomic signature predicts immunotherapy response specifically in chemotherapy-primed patients, a treatment-context specificity not shown by established immune biomarkers. These findings support prospective validation of the signature to guide chemotherapy– immunotherapy sequencing.

**Significance:** Chemotherapy is now added to immunotherapy across a growing list of cancers, yet every eligible patient receives it even though only some tumors need it, and current biomarkers (PD-L1, tumor mutational burden, microsatellite instability) cannot identify which. We report the first biomarker to predict benefit from the chemotherapy component itself: a baseline gene-expression signature that flags likely responders in chemotherapy-primed settings while remaining silent under immunotherapy alone. This treatment-context specificity shifts the biomarker question from “who responds to immunotherapy” to “who needs the chemotherapy,” pointing toward a way to spare patients unnecessary toxicity and rationally sequence chemo-immunotherapy.

## Introduction

Immune checkpoint inhibitors (ICIs) have transformed the treatment of advanced solid tumors, producing durable responses in a subset of patients across diverse cancer types [Topalian et al., 2023, Galluzzi et al., 2020]. Yet most patients do not respond durably, either failing to respond or developing resistance [Holder et al., 2024]. A key determinant of ICI efficacy is the tumor microenvironment (TME): tumors with abundant immune infiltration (“hot”) generally respond to checkpoint blockade, whereas those characterized by immune exclusion, immunosuppressive cell populations, and physical stromal barriers (“cold”) frequently fail [Holder et al., 2024, Pfirschke et al., 2016]. Importantly, preclinical studies have demonstrated that immunogenic chemotherapy can convert “cold” tumors to checkpoint-blockade-responsive “hot” states [Pfirschke et al., 2016], establishing that the TME is a modifiable, and therefore predictable, determinant of treatment outcome [Galluzzi et al., 2024].

Accumulating evidence indicates that chemotherapy can remodel the TME in ways that sensitize tumors to subsequent immunotherapy [Galluzzi et al., 2020, 2024]. Chemotherapy depletes immunosuppressive cell populations including regulatory T cells and myeloid-derived suppressor cells, and upregulates antigen presentation machinery [Galluzzi et al., 2020, 2024]. Clinically, neoadjuvant chemotherapy has been shown to increase CD8+ T cell infiltration and PD-L1 expression in esophageal squamous cell carcinoma [Liu et al., 2024], and treatment sequence is increasingly recognized as an important, though not yet definitively optimized, determinant of benefit [Galluzzi et al., 2020]. These insights have translated into landmark clinical trials: in CheckMate 816, neoadjuvant nivolumab plus chemotherapy increased the pathologic complete response rate from 2.2% to 24% in resectable NSCLC [Forde et al., 2022]; in KEYNOTE-522, neoadjuvant pembrolizumab plus chemotherapy significantly improved event-free survival in triple-negative breast cancer [Schmid et al., 2022]; and a meta-analysis of five phase III trials confirmed that perioperative chemo-immunotherapy reduces the risk of death by 32% [Zhang et al., 2024]. However, not all patients experience favorable TME changes: in esophageal cancer, only 65% showed PD-L1 upregulation after neoadjuvant chemo-immunotherapy [Liu et al., 2024], highlighting the heterogeneity of chemotherapy-induced TME conversion and the need to predict who will benefit.

Current biomarkers for ICI response are inadequate for this task. PD-L1 expression, tumor mutational burden (TMB), microsatellite instability (MSI), and even composite TME-based scores such as the Immunoscore [Galon and Bruni, 2019] provide static, single-dimensional assessments that fail to capture the dynamic and multifaceted nature of the TME [Holder et al., 2024]. In CheckMate 816, TMB was not identified as a determinant of benefit in subgroup analyses [Forde et al., 2022]. A pan-cancer meta-analysis of over thousand ICI-treated patients demonstrated that combining tumor-intrinsic and TME-based features in multivariable models significantly outperforms any single biomarker [Litchfield et al., 2021], and translational analyses from the NEOSTAR trial revealed that neoadjuvant chemo-immunotherapy induces quantifiable TME remodeling that correlates with pathologic response [Cascone et al., 2023]. Yet no existing approach specifically addresses the question of predicting which patients’ TME will convert favorably in response to chemotherapy as a means of forecasting subsequent immunotherapy benefit.

These observations suggest that the capacity for chemotherapy-induced TME conversion is an intrinsic, measurable property of the pre-treatment tumor, and that capturing this property could enable prospective patient stratification. We reasoned that by modeling the TME as a multidimensional landscape using cell type deconvolution and functional pathway scores, we could quantify treatment-induced shifts in paired pre-/post-chemotherapy biopsies, identify the subset of patients whose TME converts to a favorable state, and derive a gene-level signature of conversion propensity from pre-treatment expression alone. We then validated this signature across multiple independent cohorts of patients treated with chemo-immunotherapy or immunotherapy alone, testing the specific prediction that it would discriminate responders only among chemotherapy-primed patients.

## Results

### Identification of Chemotherapy-Induced TME Converters

To identify patients whose tumors are poised to benefit from chemotherapy prior to immunotherapy, we designed a two-stage analytical framework (**Figure 1**). In the first stage, we trained a model to estimate the favorability of a tumor’s microenvironment for immunotherapy response. In the second stage, we applied this model to paired pre-/post-chemotherapy biopsies to identify patients whose TME converted from an unfavorable to a favorable state following chemotherapy, and derived a gene signature that distinguishes these converters at baseline.

**Figure 1:**
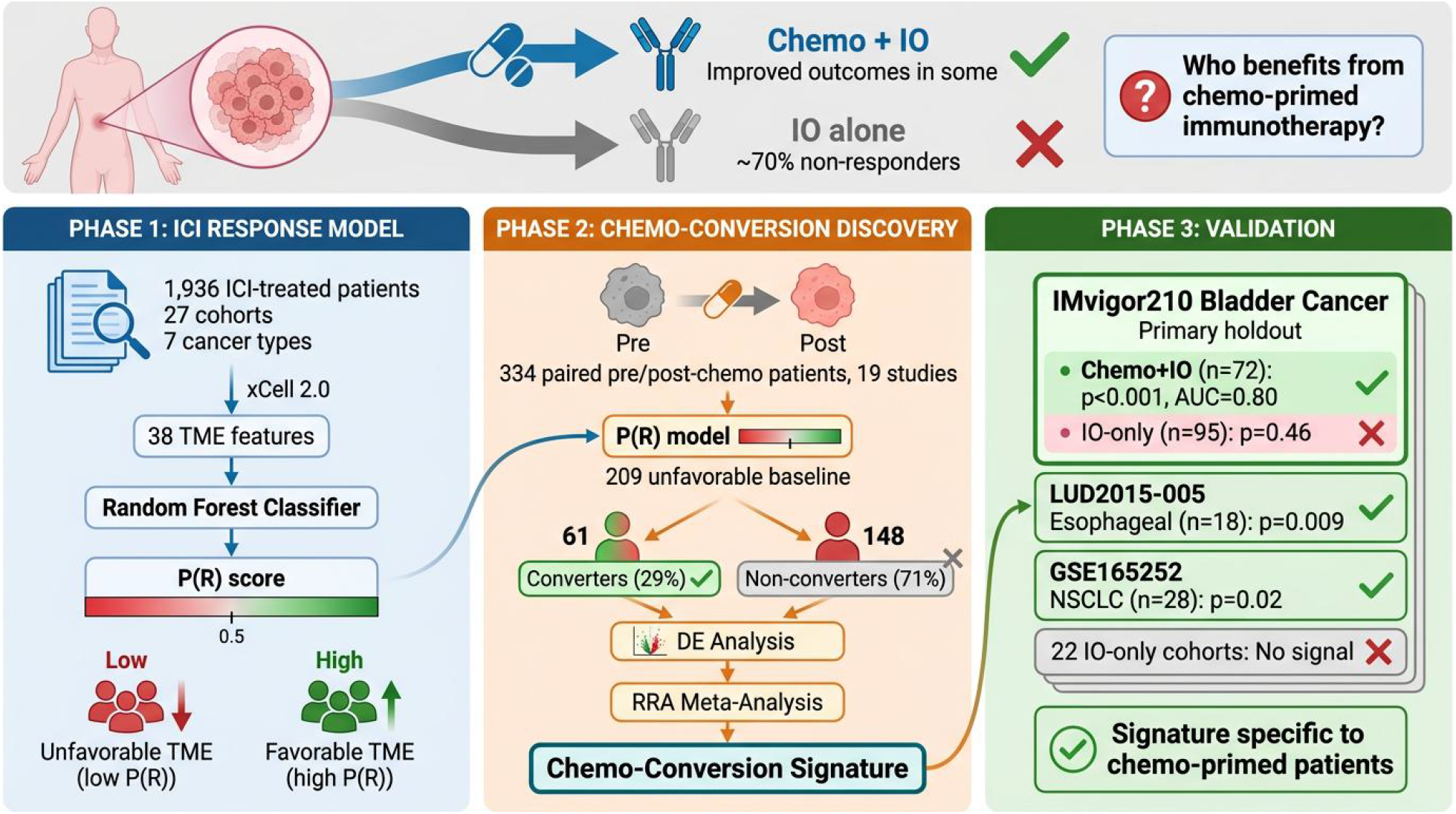
Study overview. The analytical framework comprises three phases. **Phase 1:** A random forest classifier trained on 1,936 ICI-treated patients across 27 cohorts predicts immunotherapy response probability (P(R)) from 38 xCell 2.0-derived TME features, stratifying tumors into favorable and unfavorable immune states. **Phase 2:** The model is applied to 334 paired pre-/post-chemotherapy biopsies from 19 studies; among 209 patients with unfavorable baseline TME, 61 (29%) converted to favorable status after chemotherapy. Differential expression analysis between converters and non-converters, aggregated via Robust Rank Aggregation, yields the 50-gene chemo-conversion signature. **Phase 3:** The signature is validated in independent chemo-immunotherapy cohorts (IMvigor210 bladder cancer, LUD2015-005 esophageal, GSE165252 NSCLC) and tested for specificity against 22 immunotherapy-only cohorts.

For the first stage, we assembled 1,936 ICI-treated patients across 27 cohorts and 7 cancer types (documented in [Angel et al., 2025]) and trained a random forest classifier to predict P(R), the probability of immunotherapy response, from TME features derived by xCell 2.0 [Angel et al., 2025] cell type deconvolution and functional gene expression pathway scores [Bagaev et al., 2021]. Boruta feature selection identified 38 informative TME features (Figure S1). The most predictive features included CD8+ T cell subsets (naive, exhausted, PD1-high), cancer-associated fibroblasts, gamma-delta T cells, and functional scores for checkpoint molecules and co-stimulatory ligands (Figure S2A), capturing both immune effector and stromal barrier dimensions of the TME. The model achieved a median leave-one-dataset-out AUC of 0.654 across cancer types (out-of-bag AUC = 0.639; out-of-bag calibration in Figure S2B).

For the second stage, we applied this model to paired pre- and post-chemotherapy tumor samples from 334 patients across 19 independent studies spanning RNA-Seq (119 pairs) and microarray (215 pairs) platforms (Supplementary Table S2). Of the paired patients, 209 (63%) had unfavorable baseline TME (P(R) ≤ 0.50).

Following chemotherapy, 61 of these 209 patients (29%) converted to favorable TME states, while 148 remained unfavorable (**Figure 2A**). Converters were distributed across 15 of 19 datasets, with no single dataset contributing more than 25% (top three, 48%), indicating that conversion is not a dataset-specific effect. Median P(R) among converters rose from 0.44 to 0.58 (median ΔP(R) = +0.139) versus minimal change in non-converters (−0.024). Conversion was not confined to a particular chemotherapy class, occurring comparably across anthracycline-, platinum-, and taxane/fluoropyrimidine-based regimens (between-class moderator *p* = 0.47; **Supplementary Figure S3, Supplementary Table S3**).

**Figure 2:**
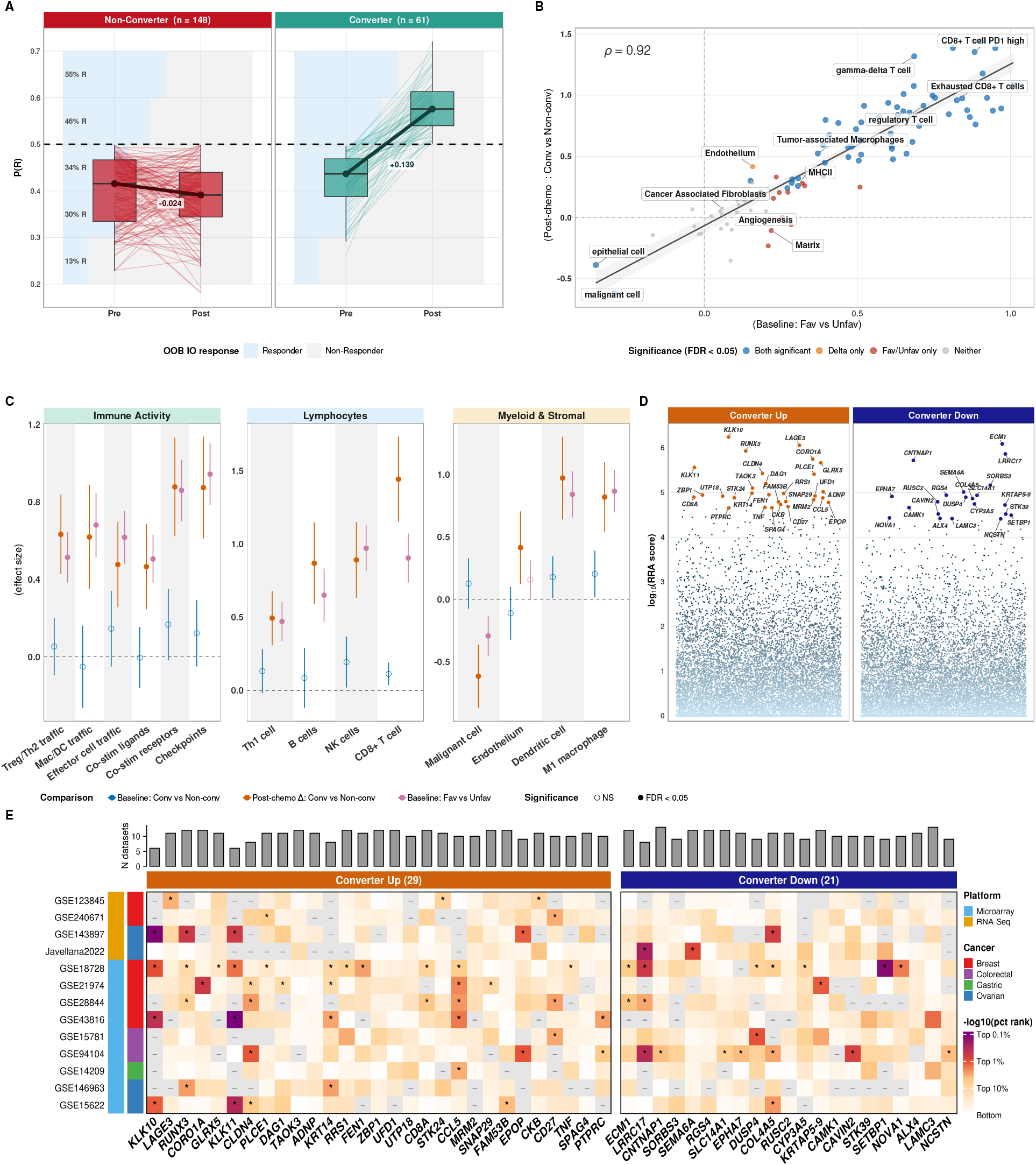
Converter identification, TME remodeling, and signature derivation. **(A)** Patient P(R) trajectories from pre-to post-chemotherapy. Dashed line: classification threshold (0.50). **(B)** TME effect size correlation: post-chemotherapy Δ (converter vs. non-converter) versus baseline favorable vs. unfavorable effect sizes across TME features (Spearman *ρ* = 0.92). **(C)** Forest plot of mixed-effect model effect sizes (95% CI) for 14 representative TME features across three comparisons: baseline converter vs. non-converter, post-chemotherapy Δ, and baseline favorable vs. unfavorable. Filled circles: FDR *<* 0.05. **(D)** RRA scores for all genes, with 50 signature genes highlighted (orange: upregulated; blue: downregulated). **(E)** Per-dataset percentile rank heatmap of signature genes across 13 datasets. Asterisks: p *<* 0.05; dashes: opposite direction.

To characterize the underlying TME changes, we compared 103 deconvolution-derived TME features between converters and non-converters using linear mixed models (dataset as random intercept) across three contrasts: baseline converter vs. non-converter, post-chemotherapy change, and baseline favorable vs. unfavorable (**Figure 2B–C**; **Supplementary Figure S4**). Strikingly, the two groups were indistinguishable at baseline (no feature reached FDR *<* 0.05; no separation by principal-component analysis; **Supplementary Figure S5**). After chemotherapy they diverged sharply: 67 of 103 features differed (FDR *<* 0.05), with converters showing broad immune activation and tumor-cell depletion, and these shifts closely mirrored the profile that distinguishes immunotherapy-favorable from unfavorable tumors at baseline (Spearman *ρ* = 0.92, *p <* 10^*−*41^). Because converters were indistinguishable at the cell-composition level, we turned to gene expression to find baseline features predictive of conversion.

### Derivation of the Chemo-Conversion Signature

To find baseline genes predictive of conversion, we performed differential expression within each of the 13 derivation datasets and aggregated the results by Robust Rank Aggregation (RRA) [Kolde et al., 2012], which identifies genes with consistent rank across cohorts despite modest per-dataset effects. Of 43,862 genes, 1,940 showed significant cross-dataset convergence (RRA score *<* 0.01); the top 50 (29 upregulated, 21 downregulated) formed the chemo-conversion signature (**Figure 2D**; Supplementary Table S4), with the median gene concordant in 79% of contributing datasets (**Figure 2E**). As an internal check, baseline signature score separated converters from non-converters in the discovery cohorts (AUC = 0.71; **Supplementary Figure S6**), as expected given that the signature was derived from these groups.

### The Chemo-Conversion Signature Predicts Response Specifically in Chemotherapy-Primed Patients

To test whether the chemo-conversion signature predicts immunotherapy response specifically in chemotherapy-primed patients, we evaluated it in IMvigor210 [Mariathasan et al., 2018] (a single-arm phase II trial of atezolizumab in metastatic urothelial carcinoma, and the only cohort containing both chemo-primed and IO-only patients, which allowed the treatment-context interaction to be tested within a single trial) and in five independent chemo-immunotherapy validation cohorts: GSE165252 (NSCLC), LUD2015-005 [Carroll et al., 2023] (esophageal adenocarcinoma), GSE207422 (NSCLC), GSE241876 (breast cancer), and the TONIC trial [Voorwerk et al., 2019] (triple-negative breast cancer). For specificity, we additionally tested 22 immunotherapy-only cohorts that were used for RF model training but excluded from signature derivation. The five validation cohorts span multiple tumor types and chemotherapy regimens and are individually small (n = 9–28), and were therefore assessed both individually and in aggregate. The signature should predict response in chemo-primed patients but not in those receiving immunotherapy without preceding chemotherapy, even when both groups share comparable baseline TME (P(R) ≤ 0.50).

In IMvigor210, among Chemo+IO patients with unfavorable baseline TME (n = 72; 13 responders, 59 non-responders), responders had significantly higher signature scores than non-responders (one-sided *p <* 0.001; AUC = 0.80, 95% CI: 0.67–0.94; **Figure 3A**). This estimate rests on only 13 responders, so the confidence interval is wide (~0.27 units) and the point estimate should be read together with that interval. In contrast, the signature showed no significant signal in the IO-only subgroup at the same unfavorable baseline (n = 95; *p* = 0.46; AUC = 0.51, 95% CI: 0.37–0.65; Hedges’ *g* = 0.01, 95% CI: −0.50 to 0.52). A logistic-regression interaction test combining both arms confirmed this pattern (interaction OR = 4.42, 95% CI: 1.53–12.73, *p* = 0.006, adjusted for baseline P(R); unadjusted OR = 3.53, 95% CI: 1.32–9.44, *p* = 0.012).

**Figure 3:**
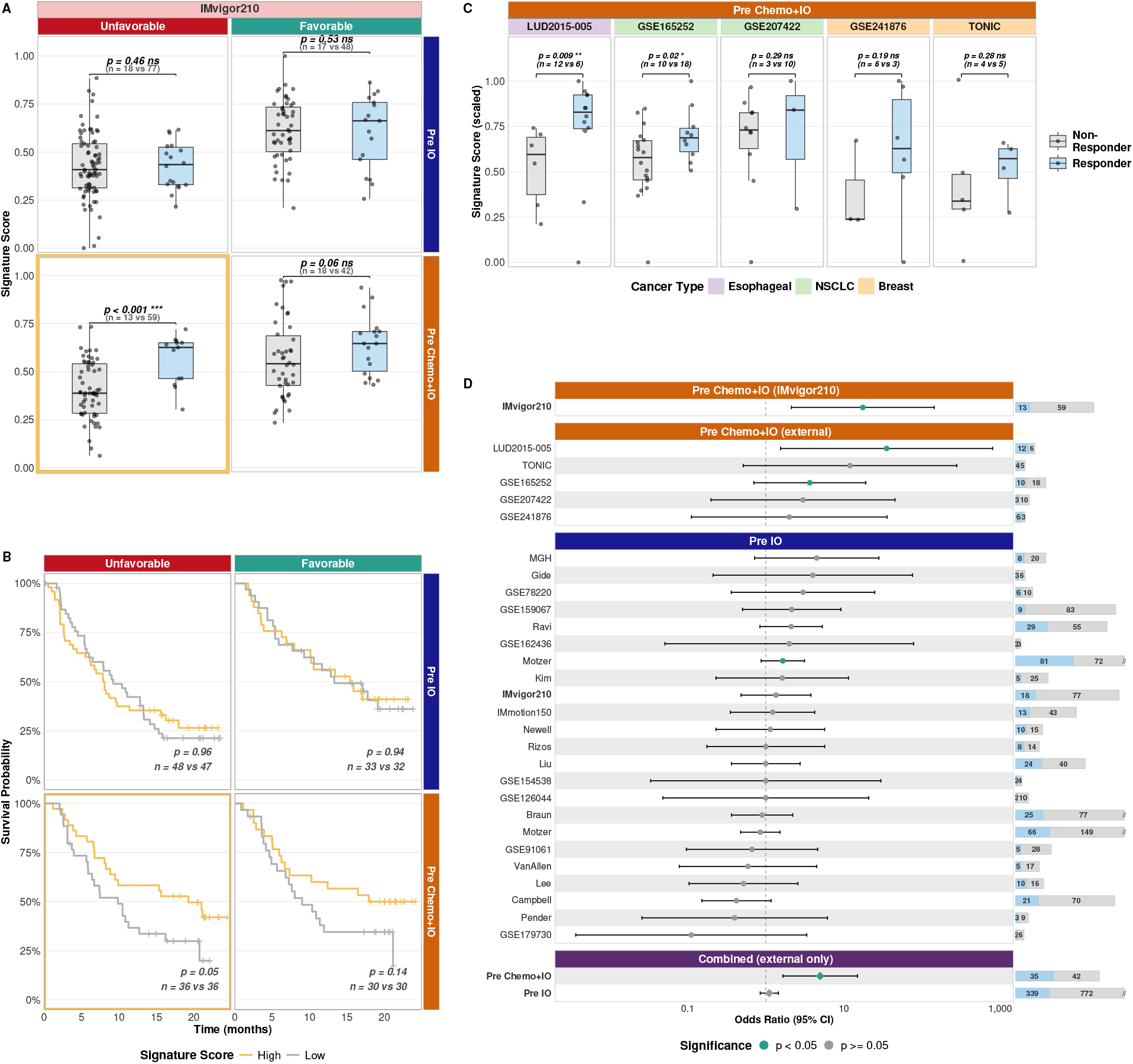
Signature validation across cohorts and treatment contexts. **(A)** Signature score distributions for responders vs. non-responders in IMvigor210, arranged in a 2×2 grid by baseline TME status and treatment context. The signature discriminates responders only in unfavorable-baseline, chemo-primed patients (highlighted panel; p *<* 0.001). P-values: one-sided Wilcoxon rank-sum tests. **(B)** Kaplan-Meier overall-survival curves in the same 2×2 layout, stratified by median signature score (high vs. low). In the chemo-primed, unfavorable-baseline panel, high score trends toward longer survival (HR = 0.55, 95% CI: 0.30–1.02, *p* = 0.056; CI crosses 1, i.e. a non-significant trend); no association is seen in the other panels. **(C)** Signature score distributions across the five chemo-immunotherapy validation cohorts (GSE165252, LUD2015-005, GSE207422, GSE241876, and TONIC), restricted to unfavorable-baseline patients. **(D)** Forest plot of odds ratios, grouped top-to-bottom into IMvigor210 (pre-chemo+IO), the five external chemo+IO validation cohorts (pre-chemo+IO), the immunotherapy-only specificity cohorts (pre-IO: the IMvigor210 IO-only arm and 22 external controls), and pooled summary estimates. To keep IMvigor210 distinct, the pooled (“combined”) estimates are computed from the external cohorts only and exclude IMvigor210. The signature predicts response in chemo-primed cohorts but shows a bounded effect in immunotherapy-only settings (pooled Hedges’ *g* = 0.08, 95% CI: −0.08 to 0.23; nominally significant in 1/22 individual cohorts).

The two arms are not randomized (**Supplementary Table S5**). The IO-only arm is a mixture of 38 chemotherapy-naïve patients and 57 biopsied after earlier platinum, and differs from the Chemo+IO arm in baseline fitness as well as chemotherapy exposure. Only ECOG performance status was significantly imbalanced, while the signature score itself was balanced across arms (*p* = 0.56), indicating that the interaction reflects a differential association with response rather than a baseline shift in score; adjusting for ECOG left it essentially unchanged (OR = 4.33, *p* = 0.007). Because prior platinum and biopsy timing are perfectly confounded with arm, we addressed them by restriction, comparing Chemo+IO only with the 38 chemo-naïve, baseline-biopsied IO-only patients, for whom subsequent chemotherapy is the only systematic difference. The interaction persisted (OR = 3.70, *p* = 0.031; ECOG-adjusted OR = 3.39, *p* = 0.048), and the chemo-naïve patients alone showed no signal (*p* = 0.36). This matched comparison is the least confounded but also the limiting evidence: with only 38 chemo-naïve controls the adjusted interaction is marginal, so the data are consistent with, but do not definitively establish, a signal specific to chemotherapy-primed patients (**Figure 3A**; see Limitations).

The signature also showed baseline-TME specificity: in IMvigor210 patients with favorable baseline TME it discriminated responders in neither arm (Chemo+IO *p* = 0.06; IO-only *p* = 0.53), consistent with chemotherapy-induced remodeling mattering only when an unfavorable immune barrier must be overcome (**Figure 3A**).

Overall survival, an exploratory and underpowered endpoint, showed a non-significant trend toward longer survival in high-signature Chemo+IO patients (median-split HR = 0.55, 95% CI: 0.30–1.02, *p* = 0.056; continuous HR = 0.75 per SD, *p* = 0.058; **Figure 3B**) and no association in the IO-only subgroup. Both intervals cross 1, so this is hypothesis-generating rather than evidence of a survival benefit.

The interaction was robust to the unfavorable-baseline threshold: across 11 cutoffs from 0.40 to 0.60, the IO-only subgroup remained non-significant throughout (AUC near 0.5) while the Chemo+IO subgroup stayed near AUC 0.8, and the interaction held wherever the Chemo+IO subgroup retained testable size (**Supplementary Figure S7**). The 0.50 threshold is the Youden-optimal cutoff of the favorability model, not a value tuned to this contrast.

We next evaluated the signature in the five independent chemo-immunotherapy validation cohorts spanning NSCLC, esophageal adenocarcinoma, and breast cancer (GSE165252, LUD2015-005, GSE207422, GSE241876, and the TONIC trial). In all five, responders had higher baseline signature scores than non-responders (**Figure 3C**). The association was individually significant in the two larger cohorts (GSE165252: one-sided *p* = 0.02, AUC = 0.73, 95% CI: 0.55–0.92; LUD2015-005: *p* = 0.009, AUC = 0.85, 95% CI: 0.65–1.00) and directionally consistent, though not individually significant, in the three smaller cohorts (n = 9–13). Because each cohort is small, we combined them by random-effects meta-analysis, which confirmed a significant overall association between baseline signature score and chemo-immunotherapy response (pooled Hedges’ *g* = 0.59, 95% CI: 0.11–1.08, *p* = 0.017).

To rule out that the signature merely predicts general immunotherapy response, we applied it to 22 IO-only cohorts restricted to unfavorable baseline TME. A random-effects meta-analysis gave a small, non-significant pooled effect (Hedges’ *g* = 0.08, 95% CI: −0.08 to 0.23; *p* = 0.35; low heterogeneity, no funnel-plot asymmetry). Equivalence testing (TOST) [Lakens, 2017] against a |*g*| ≤ 0.20 margin gave *p* = 0.061, narrowly short of formal equivalence; the upper bound (*g* = 0.23) nonetheless constrains any IO-only effect to at most a small magnitude, and the signature reached nominal significance in only 1 of 22 cohorts (**Figure 3D**). Any residual IO-only association is therefore, at most, clinically marginal and substantially weaker than in the chemotherapy-primed setting.

### Functional Characterization Reveals Four Interconnected Biological Programs in Converter Tumors

To characterize the biological processes distinguishing converter from non-converter tumors at base-line, we performed a meta-analytic pathway enrichment analysis following the MAPE-G framework [Shen and Tseng, 2010]. In our implementation, a per-gene random-effects meta-analysis across all 13 paired chemotherapy datasets produced a single consensus ranked gene list (19,217 genes), which was then subjected to GSEA against 6,631 Hallmark, GO Biological Process, and Reactome pathway gene sets, yielding 1,077 significantly enriched pathways (FDR *<* 0.05). After deduplication to remove redundant gene sets, 27 representative pathways emerged, organized into four functional modules (**Figure 4A–B**).

**Figure 4:**
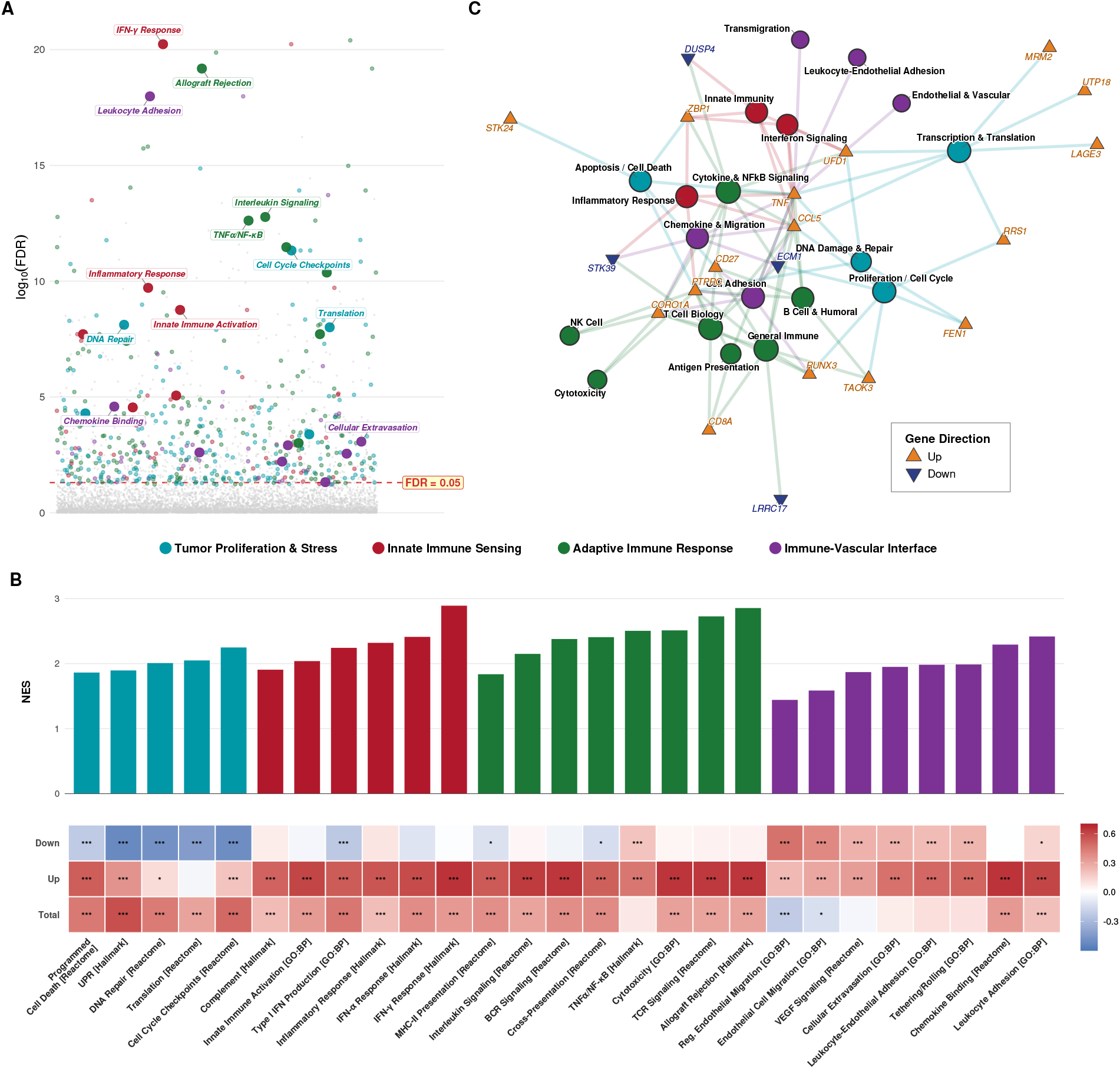
Functional characterization of the chemo-conversion signature. **(A)** Enrichment landscape showing − log_10_(FDR) for 6,631 pathways tested. The 27 curated representative pathways are shown as enlarged points colored by module; for legibility, the most significant pathways per module are labeled, with the complete set named in panel (B). Red dashed line: FDR = 0.05. **(B)** NES barplot of 27 curated pathways organized by four functional modules, with pathway score correlation heatmap (bottom) showing Spearman *ρ* between each pathway and the signature components across all response-evaluable IMvigor210 patients (n = 298; this descriptive correlation is not restricted to the unfavorable-baseline subset). **(C)** Gene-submodule membership map linking 20 signature genes (triangles) to the 20 biological submodules (circles, colored by module) whose gene sets contain them. Genes such as TNF, CCL5, and PTPRC are members of submodules spanning multiple modules.

The 27 enriched pathways organized into four modules (**Figure 4A–B**). Converter tumors were enriched for proliferation and stress programs (E2F and MYC targets, G2M checkpoint, DNA replication) coupled with proteotoxic stress and apoptotic priming, with partial-correlation analysis confirming the unfolded-protein-response and apoptotic signals as independent of proliferation, consistent with proliferating, stress-loaded cells primed for death on chemotherapy exposure. They were also enriched for innate immune sensing, led by IFN-*γ* response (the strongest enrichment in the dataset), with IFN-*α* response, complement, and inflammatory programs.

Downstream, converter tumors showed adaptive immune effector programs (allograft rejection, T- and B-cell-receptor signaling, antigen cross-presentation, leukocyte-mediated cytotoxicity, TNF*α*/NF-*κ*B); the innate and adaptive programs were tightly correlated (pairwise *ρ >* 0.85), forming a single coordinated immune axis. A fourth module captured the immune-vascular interface, with the leukocyte-extravasation cascade and VEGF signaling enriched and endothelial migration (but not vasculogenesis) activated, while TGF-*β*, which drives endothelial anergy and immune exclusion [Griffioen et al., 1996, Mariathasan et al., 2018], was anti-correlated with the signature, consistent with functional rather than anergic vasculature.

These modules cohere into one program. Across response-evaluable IMvigor210 patients, immune and proliferative pathway scores correlated strongly with the upregulated signature component and vascular pathways intermediately (**Figure 4B**), and several signature genes (notably TNF, CCL5, PTPRC, and CD27) were shared across submodules spanning multiple modules (**Figure 4C**), indicating that converter tumors harbor an integrated state linking tumor vulnerability, immune readiness, and vascular access rather than independent axes.

## Discussion

Biomarkers for immunotherapy response have almost uniformly measured one static property of the pre-treatment tumor: how inflamed it already is, whether through PD-L1 expression, tumor mutational burden, microsatellite instability, or composite microenvironment scores [Holder et al., 2024]. The decision facing a clinician who is considering chemo-immunotherapy is a different one. It is not whether a tumor is hot today, but whether it will become hot if chemotherapy is given first, and therefore whether the chemotherapy component is worth its toxicity. The chemo-conversion signature reframes the biomarker question along exactly this line. Rather than scoring a tumor’s current immune state, it predicts a treatment-induced transition, flagging tumors poised to convert from an unfavorable to a favorable microenvironment after chemotherapy. It is thus a predictive biomarker of chemotherapy benefit rather than a prognostic marker of immunotherapy response, a distinction that, to our knowledge, has not previously been demonstrated for immune-oncology treatment sequencing.

The strongest evidence that this is a different kind of biomarker is its treatment-context specificity. Within IMvigor210 the signature separated responders from non-responders among chemotherapy-primed patients but not among patients given immunotherapy without preceding chemotherapy, despite comparable baseline TME; the same direction held across five independent chemo-immunotherapy cohorts, and a meta-analysis of 22 immunotherapy-only cohorts bounded any effect there to a small magnitude. Tellingly, none of five established immune-activation signatures (IFN-*γ*, the Tumor Inflammation Signature [Ayers et al., 2017], the Gene Expression Profile, a T-effector signature, and IPRES) reproduced the interaction (**Supplementary Figure S8, Supplementary Table S6**): a static measure of immune infiltration cannot, by construction, distinguish a tumor that needs chemotherapy from one that does not, whereas this signature can. These conclusions rest on modest numbers (13 responders in the primary analysis, a borderline replication) and need larger confirmation, but the consistent direction across cohorts and the failure of canonical signatures to reproduce the effect argue that it is real.

How the signature achieves this points to a second, less expected result. At baseline, converter and non-converter tumors were essentially indistinguishable by cell composition: none of 103 deconvolution-derived microenvironment features separated the two groups, and principal-component analysis showed no separation along any major axis. The capacity to convert was nonetheless legible at the level of gene expression, where a compact transcriptional program identified future converters before any treatment. Conversion propensity, in other words, is not captured by how much immune infiltrate a tumor already carries; it is a latent, coordinated molecular state that conventional cell-type quantification does not see. This is why the signature correlates only weakly with immune-hot scores (**Supplementary Figure S8**) yet predicts a response those scores miss.

Pathway analysis suggests what that latent state is made of: one coherent picture of a tumor primed to convert. Converter tumors were enriched for proliferating, stress-loaded programs that poise cancer cells for immunogenic cell death, set within a microenvironment already equipped with innate sensing machinery and adaptive effectors and served by a functional vasculature able to deliver immune cells to the site. In this model chemotherapy is the trigger that releases antigens and danger signals from cells on the edge of immunogenic death, and subsequent checkpoint blockade sustains the response [Pfirschke et al., 2016, Galluzzi et al., 2024]. This account is a hypothesis built from observational enrichment data and preclinical precedent for immunogenic chemotherapy [Pfirschke et al., 2016], and will require direct experimental test.

The clinical promise is straightforward: to spare patients whose tumors will not convert the toxicity and cost of a chemotherapy component they do not need, a stake illustrated by trials such as KEYNOTE-859, in which PD-L1-negative gastric-cancer patients gained no survival benefit from added immunotherapy while still being exposed to its added treatment-related toxicity [Rha et al., 2023]. The most immediate applications are in sequential settings: identifying, before treatment, patients whose tumors are likely to be converted by chemotherapy and thus to benefit from subsequent immunotherapy, and enriching trials that compare sequential chemo-immunotherapy with immunotherapy alone. Extension to concurrent regimens (KEYNOTE-189, KEYNOTE-355, CheckMate 649), most readily via retrospective analysis of biomarker-stratified trial subsets, together with a standardized assay and prospective validation, are the steps toward clinical use.

Several limitations apply. First, the central interaction rests on a non-randomized comparison of IMvigor210 subgroups that differ in biopsy timing and baseline fitness as well as chemotherapy; we mitigated this by restriction to a matched comparison, ECOG adjustment, and corroboration across 22 immunotherapy-only cohorts, but residual confounding cannot be excluded and definitive proof requires a randomized trial of chemo-immunotherapy versus immunotherapy alone. Second, the supporting cohorts are small (13 responders in the primary analysis, a statistically borderline replication, and the smaller validation cohorts, which are individually underpowered), so the estimates are imprecise; moreover, the discovery cancer types (breast, ovarian, colorectal, gastric, sarcoma) do not overlap the primary or replication validation cohorts, so the signature has not been confirmed in the cancer types from which it was derived. Third, because chemotherapy can reduce tumor purity, part of the converter signal could in principle reflect cellularity change; that the signature predicts from pre-treatment biopsies and is specific to chemotherapy-primed patients argues against this, but a purity-adjusted re-derivation remains for future work. Finally, the signature was validated only where the baseline biopsy precedes chemotherapy, not in modern concurrent chemo-immunotherapy regimens, and prospective validation in a randomized trial is required before clinical use.

More broadly, our results suggest that treatment-induced changes in the tumor microenvironment are not arbitrary but are, at least in part, encoded in the pre-treatment tumor and therefore predictable. If that holds, the static hot-versus-cold framing that dominates immune-oncology biomarkers could be complemented by a class of conversion biomarkers that forecast how a given therapy will reshape a tumor’s microenvironment, and that inform not only whether to treat but in what order. The chemo-conversion signature is a first concrete instance of this idea, and a step toward matching each patient to the treatment sequence their tumor is actually poised to benefit from.

## Methods

### Data Collection and Preprocessing

We assembled three transcriptomic collections from public repositories (Gene Expression Omnibus, Synapse, published data): (i) an ICI training set of 1,936 pre-treatment samples from 27 cohorts across 7 cancer types and multiple regimens (documented in [Angel et al., 2025]); (ii) a paired pre-/post-chemotherapy discovery set (Supplementary Table S2); and (iii) six validation cohorts reserved for signature testing. All validation cohorts were withheld from model training, converter identification, and signature derivation.

Raw data were processed in their native format (RNA-seq as TPM; Affymetrix microarrays by RMA with updated Brainarray CDFs) and stored with harmonized annotations. Response was defined as complete or partial response (responder) versus stable or progressive disease (non-responder) per RECIST v1.1 or study-specific criteria; replicate biopsies were collapsed to a single sample per timepoint to avoid leakage (Supplementary Methods).

### TME Feature Quantification

TME composition was quantified with xCell 2.0 [Angel et al., 2025] against four deconvolution reference panels (Blueprint/ENCODE, LM22/CIBERSORT [Newman et al., 2015], Kassandra Tumor, and a pan-cancer single-cell atlas) and complemented by 29 functional gene-expression (FGE) pathway scores [Bagaev et al., 2021], yielding 144 features per sample (per-source counts in Supplementary Table S1).

### Batch Effect Correction via Per-Dataset Normalization

Batch effects across studies and platforms were removed by per-dataset median-MAD normalization (values winsorized to the 1st–99th percentiles, z-scores capped at ±5), retaining 134 features after a near-zero-MAD filter; each validation cohort was normalized using its own statistics to prevent leakage (formula in Supplementary Methods).

### Immunotherapy Response Model Development

We trained a random forest (ranger [Wright and Ziegler, 2017]) to predict P(R) from the normalized TME features, optimizing hyperparameters by cancer-type-balanced leave-one-dataset-out (LODO) cross-validation over five cancer-type folds, each anchored by the largest cohort of that type (Melanoma [Newell et al., 2022], RCC [Motzer et al., 2020], NSCLC [Ravi et al., 2023], HNSCC [Foy et al., 2022], Gastric [Kim et al., 2018]); the selected model reached a median LODO AUC of 0.654 (Supplementary Tables S7–S8).

Feature selection used the Boruta algorithm [Kursa and Rudnicki, 2010], which retains features whose importance significantly exceeds that of randomly permuted shadow features. This selected 38 informative features from the 134, outperforming nine alternative selection strategies in out-of-bag AUC; the source composition of the retained features is shown in Figure S1. Feature selection was performed on the full training set rather than nested within each leave-one-dataset-out fold; the close agreement between out-of-bag and LODO AUC (0.639 vs. 0.654) indicates that any resulting leakage is minimal, and the rationale and implications of this design choice are detailed in Supplementary Methods.

The final model was trained on all 1,936 samples (38 features, inverse-frequency class weights) and applied to every discovery and validation cohort, each normalized independently; unfavorable baseline TME was defined as P(R) ≤ 0.50 (near the out-of-bag Youden-optimal cutoff of 0.470).

### Identification of Chemotherapy-Induced TME Converters

We assembled 19 studies with matched pre-/post-chemotherapy biopsies (334 patients; platinum-, taxane-, and combination regimens; Supplementary Table S2). Because these cohorts were excluded from RF training, their P(R) predictions are out-of-cohort. A patient was a converter if pre-chemotherapy P(R) was ≤ 0.50 and rose above it after treatment, and a non-converter otherwise; conversion rates were comparable across platforms (Supplementary Methods).

### Derivation of the Chemo-Conversion Signature

Differential expression was computed within each paired dataset with limma [Ritchie et al., 2015] (RNA-seq log_2_(TPM+1)-transformed; microarray used directly), and per-dataset results were aggregated by Robust Rank Aggregation (RRA) [Kolde et al., 2012], which scores genes by rank consistency across cohorts. Genes with RRA score *<* 0.01 were candidates, and the top 50 (29 upregulated from 1,127, 21 downregulated from 813) formed the signature (Supplementary Table S4); overlap with the TME-feature marker panels is reported in Supplementary Table S9.

### Signature Scoring and Validation

Per-sample scores were computed with singscore [Foroutan et al., 2018] as the normalized mean rank of the upregulated genes combined with the reverse-ranked downregulated genes, a rank-based metric robust to cross-platform differences; association with response was tested by one-sided Wilcoxon rank-sum test (alternative: responders *>* non-responders), with two-sided values reported in Supplementary Table S10.

To summarize results across the five chemo-immunotherapy validation cohorts, per-cohort effect sizes (Hedges’ *g*) were combined by random-effects (REML) meta-analysis (`metafor::rma()`).

Within IMvigor210, biopsy timing relative to platinum defined two subgroups, Chemo+IO and IO-only (the latter a mixture of chemotherapy-naïve and post-platinum-biopsy patients); after excluding unclassifiable and response-missing cases, 72 Chemo+IO and 95 IO-only patients remained at unfavorable baseline. The full construction and counts are given in Supplementary Methods.

The signature×arm interaction was tested by logistic regression (response ~ signature score × arm + baseline P(R)) on standardized scores in the unfavorable-baseline population. Because the arms are non-randomized, the model was adjusted for ECOG performance status (the one imbalanced covariate; Supplementary Table S5), and prior-platinum/biopsy-timing confounding, which cannot be adjusted by regression, was addressed by restriction to a chemotherapy- and biopsy-timing-matched comparison (Chemo+IO versus chemo-naive IO-only); residual confounding is discussed in the Limitations.

The signature was evaluated in six chemo-immunotherapy validation cohorts restricted to unfavorable baseline TME: **IMvigor210** (urothelial; the only cohort with both arms), **GSE165252** (NSCLC), **GSE207422** (NSCLC), **GSE241876** (breast), the **TONIC trial** [Voorwerk et al., 2019] (triple-negative breast), and the IO-then-chemo cohort **LUD2015-005** (esophageal); detailed per-cohort descriptions are in Supplementary Methods. Overall survival in IMvigor210 was compared by median-split log-rank test with Cox-model hazard ratios.

To confirm chemotherapy specificity, the signature was tested across 22 immunotherapy-only training cohorts (each restricted to unfavorable-baseline patients with at least three responders and non-responders), using out-of-bag predictions for stratification to avoid overfitting. Per-cohort effect sizes (Hedges’ *g*) were combined by the same random-effects (REML) meta-analysis used for the validation cohorts. Because non-significance does not establish absence of effect, we additionally performed equivalence testing (two one-sided tests) against a margin of |*g*| ≤ 0.20 (Cohen’s small-effect threshold, ΔAUC ≈ 0.05) [Lakens, 2017]; the full procedure is described in Supplementary Methods.

Biological processes distinguishing converters from non-converters at baseline were characterized by meta-analytic pathway enrichment (following the MAPE-G framework [Shen and Tseng, 2010]): in our implementation, a per-gene random-effects meta-analysis across the 13 datasets produced a consensus ranked list (19,217 genes), tested by GSEA against 6,631 Hallmark, GO Biological Process, and Reactome gene sets, yielding 1,077 enriched pathways (FDR *<* 0.05).

Enriched pathways were deduplicated and assigned to biological submodules, yielding 27 representative pathways across four modules (tumor proliferation and stress response, innate immune sensing, adaptive immune response, and immune-vascular interface). The deduplication criteria and the independence analyses used to identify distinct biological axes are detailed in Supplementary Methods.

### Statistical Analysis

All analyses used R 4.3 with a significance threshold of *α* = 0.05. Effect sizes were quantified as AUC (pROC [Robin et al., 2011]) and Hedges’ *g*; Benjamini-Hochberg FDR correction was applied to RRA, pathway enrichment, and TME-feature comparisons; and linear mixed-effect models were fitted with lmerTest [Kuznetsova et al., 2017] using Satterthwaite’s degrees-of-freedom approximation.

As a tumor-marker study that both develops a machine-learning clinical prediction model and evaluates a prognostic/predictive biomarker, this work is reported in accordance with the REMARK (biomarker) and TRIPOD+AI (machine-learning prediction-model) reporting guidelines; completed checklists are provided as **Supplementary Table S14** (REMARK) and **Supplementary Table S15** (TRIPOD+AI).

## Data Availability

All code for data processing, model training, signature derivation, and validation is available at https://github.com/AlmogAngel/chemo-immuno-dynamics and will be archived on Zenodo, with the DOI provided upon publication. Raw expression data for public datasets are available through the Gene Expression Omnibus (GEO) and Synapse using the accession numbers provided in Supplementary Tables S1 and S2. The IMvigor210 dataset is available through the IMvigor210CoreBiologies R package. The LUD2015-005 data were obtained from Carroll et al. [2023]. Processed data objects, including the trained RF model, signature gene lists, and validation results, will be deposited in a public repository upon publication. TONIC RNA-seq data are under controlled access at the European Genome-phenome Archive (accession EGAD00001004858) and require Data Access Committee approval; with approval, raw FASTQs were processed with STAR (GENCODE v32) and TPM-normalized, and the TONIC result can be reproduced by setting the `CHEMOIO_TONIC_DIR` environment variable and re-running `pipeline/07_validate_replication.R`.

## Acknowledgments

We thank members of the Aran lab for helpful discussions. This work was supported by the Israel Science Foundation (ISF), grant No. 1543/21.

