## Supplemental Material for "Predicting Immunotherapy Response Through Chemotherapy-Induced Tumor Microenvironment Remodeling"

**3 Supplementary Figures** **9**

**4 Supplementary Tables** **16**

**5 Reporting Guideline Checklists** **23**

### 1 Supplementary Methods

#### 1.1 Detailed Dataset Descriptions

##### 1.1.1 ICI Training Cohorts

Twenty-seven independent ICI-treated cohorts (1,936 samples) were used for random forest model training (Supplementary Table S1). Datasets were collected from the Gene Expression Omnibus (GEO), Synapse, and published supplementary materials. All cohorts used pre-treatment tumor biopsies (RNA-Seq, TPM-normalized). Cohort-specific response definitions (RECIST v1.1 or study-reported) were harmonized to a binary responder/non-responder classification. Response definitions were: responders versus non-responders (binary classification) for all cohorts.

##### 1.1.2 Paired Chemotherapy Cohorts

Nineteen independent studies with matched pre- and post-chemotherapy biopsies were assembled (Supplementary Table S2). Datasets span RNA-Seq and Affymetrix microarray platforms. RNA-Seq datasets: TPM-normalized. Microarray datasets: RMA-normalized with Brainarray custom CDF packages for gene-level summarization. All datasets link pre/post samples by a unique patient identifier (Pair_Key).

Eligible chemotherapy regimens included platinum-based, taxane-based, DNA-disrupting, and combination regimens (classified as Chemotherapy or Chemoradiotherapy). Datasets with chemotherapy followed immediately by immunotherapy (classified as "Chemotherapy >> Immunotherapy") were excluded from converter discovery to avoid confounding by immunotherapy-induced TME changes. Cancer types: Breast (9 datasets), Ovarian (6 datasets), Colorectal (2 datasets), Gastric (1 dataset), Sarcoma (1 dataset).

##### 1.1.3 Validation Cohorts

**IMvigor210.** Urothelial/bladder cancer cohort treated with atezolizumab (anti-PD-L1), available through the IMvigor210CoreBiologies R package. N $\approx$ 348 baseline samples. Two clinical arms were distinguished by prior treatment history:

- **Chemo+IO arm**: patients with prior platinum-based chemotherapy followed by atezolizumab (Cohort 2; $n = 72$ unfavorable-baseline).
- **IO-only arm**: patients receiving atezolizumab without immediately preceding chemotherapy (Cohort 1; $n = 95$ unfavorable-baseline). This arm is a mixture of cisplatin-ineligible

chemo-naïve patients (biopsied before any chemotherapy) and patients who received earlier-line platinum but were biopsied after it (Supplementary Table S5).

Arm assignment was based on biopsy timing relative to platinum (IMvigor210CoreBiologies annotations `Received platinum` and `Sample collected pre-platinum`). Both arms were restricted to pre-treatment samples with unfavorable baseline TME ($P(R) \leq 0.50$); patients who received platinum but whose biopsy timing relative to it was unrecorded were excluded as unclassifiable. Because prior platinum exposure and biopsy timing are perfectly confounded with arm (every Chemo+IO patient received platinum and was biopsied before it), they cannot be adjusted by regression and are instead addressed by restriction to a chemotherapy- and biopsy-timing-matched comparison in the main text.

**LUD2015-005 (supplementary).** Esophageal adenocarcinoma cohort. Patients received immunotherapy first (anti-PD-1) followed by the addition of chemotherapy. "On-treatment" biopsies were taken after the initial immunotherapy cycle but before chemotherapy addition. $N = 18$ unfavorable-baseline on-treatment samples (12 responders, 6 non-responders). Dataset obtained from Carroll et al. [2023]. Although the signature was significant in this cohort ($p = 0.009$), it is reported as a supplementary validation cohort rather than alongside the others because its on-treatment biopsies reflect an IO-then-chemo sequence (biopsy taken under anti-PD-1 pressure, before chemotherapy), which does not match the chemotherapy-primed premise the signature was derived to test.

**GSE165252.** NSCLC cohort treated with concurrent chemoradiotherapy plus anti-PD-L1. Pre-treatment samples, $N = 28$ unfavorable-baseline (10 responders, 18 non-responders). Response stored in `geo_meta[sample_id, "response:ch1"]`.

**GSE207422.** NSCLC cohort, $N = 13$ (3R/10NR), anti-PD-1 + chemotherapy, pre-treatment only (supplementary cohort).

**GSE241876.** Breast cancer cohort, $N = 9$ (6R/3NR), chemo + anti-PD-1, pre-treatment only (supplementary cohort).

**TONIC trial (restricted-access).** Triple-negative breast cancer cohort [Voorwerk et al., 2019] in which patients received a chemotherapy induction (cisplatin, doxorubicin, cyclophosphamide, or no induction control) followed by nivolumab. Pre-induction baseline biopsies were used. RNA-seq data are deposited under *controlled access* at the European Genome-phenome Archive (accession `EGAD00001004858`) and require Data Access Committee (DAC) approval;

raw and patient-level processed data are therefore *not redistributed* in this deposit. With DAC approval, raw FASTQs were aligned with STAR (GENCODE v32) and TPM-normalized, and the standard signature scoring + one-sided Wilcoxon pipeline was applied to unfavorable-baseline (P(R) $\leq$ 0.50) pre-induction biopsies in the chemotherapy-induction arms. The TONIC "Chemo+IO" subset comprises patients in the cisplatin, cyclophosphamide, and doxorubicin induction arms; the no-induction and irradiation-only arms (38 patients) were excluded as they do not deliver chemotherapy. After restriction to unfavorable baseline, the analyzable cohort is N = 9 (4R/5NR; cisplatin = 2, cyclophosphamide = 2, doxorubicin = 5). To reproduce locally, set the `CHEMOIO_TONIC_DIR` environment variable to the directory containing the processed RDS and re-run `pipeline/07_validate_replication.R`; the loader will emit an informative message (rather than silently skipping) if the data are not found.

#### 1.2 TME Feature Details

A total of 144 TME features were generated per sample:

**Table S1: TME feature sources and counts.** Per-source feature counts; full list as CSV (`tables/supplementary/stable1_tme_features.csv`).

| Source ID | Description | N Features |
|---|---|---|
| bp | Blueprint/ENCODE epigenomic profiles | ∼30 |
| lm22 | CIBERSORT LM22 signature matrix | 22 |
| kass_tumor | Kassandra tumor-associated profiles | ∼32 |
| sc_pan_cancer | Pan-cancer single-cell atlas | ∼31 |
| fge | Functional gene expression pathways | 29 |
| **Total** | | **144** |

The 29 FGE pathway scores represent the following TME processes: angiogenesis, TGF-$\beta$ signaling, effector cell trafficking, antigen presentation, immune checkpoint activity, MDSC activity, Treg enrichment, cytolytic activity, IFN-$\gamma$ signaling, and related functional axes.

After per-dataset median-MAD normalization, 134 features passed the near-zero MAD filter and were used for Boruta feature selection. Boruta identified 38 confirmed-relevant features; 96 features were rejected as non-informative.

#### 1.3 Singscore Implementation Details

Signature scores were computed using the singscore package (v1.16) in R. The implementation:

1. Rank all genes in each sample by expression level using `rankGenes()`.

2. Score the upregulated gene set (`simpleScore(upSet = up_genes)`).

3. Score the downregulated gene set (`simpleScore(upSet = down_genes, knownDirection = FALSE)`), then negate the total score.

4. Compute total score as the mean of up- and down-scores.

**Important implementation note**: When subsetting a ranked matrix produced by `rankGenes()`, the `stable` attribute must be explicitly preserved; otherwise `simpleScore()` fails with "argument is of length zero." All subsetting operations in this pipeline use a custom `subset_ranked()` wrapper that restores this attribute.

#### 1.4 Robust Rank Aggregation Details

RRA was run using the RobustRankAggreg R package (v1.2). Parameters:

- **Gene ranking within each dataset**: by p-value (ascending).
- **Minimum datasets for inclusion**: genes not detected in any dataset for a given direction were excluded from that RRA analysis.
- **Gene universe ($N$)**: number of unique genes across all 13 contributing datasets in each direction.
- **Score threshold**: genes with RRA score $< 0.01$ were considered significant candidates; the top 50 by RRA score (proportionally split: 29 upregulated from 1,127 candidates, 21 downregulated from 813 candidates) were selected for the final signature.

Thirteen datasets contributed to the RRA analysis (datasets with $\geq 2$ samples per group in both converter and non-converter classes). The remaining datasets were excluded due to insufficient sample size.

#### 1.5 Preprocessing and Additional Parameters

Raw expression matrices were stored as Bioconductor SummarizedExperiment objects with harmonized annotations (treatment timepoint, drug category, treatment strategy, clinical response); response was defined as responders (complete or partial response) versus non-responders (stable or progressive disease) per RECIST v1.1 or study-specific criteria, and replicate biopsies at the same timepoint were collapsed to a single sample to avoid information leakage. Batch effects across studies and platforms were removed by per-dataset median–MAD normalization, $z_{f,d,s} = (x_{f,d,s} - \text{median}_d(f))/\text{MAD}_d(f)$, with raw values winsorized to the

1st–99th percentiles and resulting $z$-scores capped at ±5; 134 of the 144 features survived a near-zero-MAD filter (MAD $< 0.001$), and each validation cohort was normalized using its own within-cohort statistics. Converter classification was comparable across platforms (RNA-Seq 25.8% vs. microarray 32.1%; Fisher's exact $p = 0.43$). For the pathway meta-analysis, GSEA was run against Hallmark, GO Biological Process, and Reactome gene sets of size 15–500.

##### 1.6 Feature Selection and Cross-Validation Leakage

Boruta feature selection was performed on the full 1,936-sample training set rather than nested within each leave-one-dataset-out (LODO) fold. This design permits each LODO fold to be evaluated on a feature set that was informed, at the Boruta selection step, by samples from the held-out cancer type, which could in principle inflate the LODO AUC. Two observations indicate that any such leakage is minimal. First, the out-of-bag AUC (0.639), computed without reference to the held-out folds, agrees closely with the LODO AUC (0.654), a difference of only 1.5 percentage points. Second, a fully nested Boruta-within-LODO design, which would more strictly upper-bound the leakage, is rarely applied in biomarker-discovery studies of this scale owing to its substantial computational cost (five additional Boruta runs per evaluation). We therefore interpret the reported LODO AUC as a near-honest, though not strictly nested, estimate of out-of-cancer-type generalization.

##### 1.7 Pathway Module Curation

The 1,077 significantly enriched pathways (FDR $< 0.05$) were deduplicated by assigning pathways to biological submodules and retaining, among highly correlated pathways ($\rho > 0.85$, computed across all response-evaluable IMvigor210 patients, $n = 298$), only the pathway with the strongest normalized enrichment score. Stepwise $R^2$ analysis and partial correlation controlling for the E2F proliferation signal were used to identify independent biological axes. This yielded 27 representative pathways across four modules: tumor proliferation and stress response (5), innate immune sensing (6), adaptive immune response (8), and immune-vascular interface (8). Gene-to-submodule connections were determined by pathway membership.

#### 2 Supplementary Results

##### 2.1 RF Model Performance

The random forest model trained on 38 Boruta-selected TME features achieved:

- Out-of-bag (OOB) AUC: 0.639
- Optimal classification threshold (Youden J statistic): 0.470
- Cancer-type-balanced LODO median AUC: 0.654

The OOB AUC reflects generalizability across the 1,936-sample training set spanning five cancer types. The LODO evaluation confirms that the model generalizes across tumor biology, with held-out cancer types evaluated independently.

#### 2.2 Threshold Sensitivity Analysis

To assess robustness to the choice of the unfavorable-baseline P(R) threshold, the IMvigor210 interaction analysis was repeated across thresholds spanning 0.40 to 0.60. The interaction pattern (Chemo+IO significant AND IO-only non-significant) was maintained at the majority of thresholds tested. At the two most stringent thresholds (P(R) $\leq$ 0.40 and 0.42), the Chemo+IO subgroup contained too few patients for adequate statistical power. The IO-only subgroup remained non-significant across all thresholds tested (0/all thresholds), confirming that the null result is not an artifact of threshold selection (Supplementary Figure S7).

#### 2.3 Equivalence Testing of the IO-Only Specificity Claim

A non-significant Wilcoxon test alone is consistent with both a true null effect and a small-but-real effect that the sample is underpowered to detect. To formally bound the maximum effect size compatible with the data, we performed two one-sided tests (TOST) against an equivalence margin of |Hedges' $g$| $\leq 0.20$ (Cohen's small-effect threshold, equivalent to $\Delta$AUC $\approx 0.05$). For each estimate, we tested simultaneously: (a) $g > -0.20$ and (b) $g < 0.20$; the equivalence $p$-value is the maximum of the two one-sided $p$-values, and equivalence is concluded when both nulls are rejected at $\alpha = 0.05$.

In IMvigor210 IO-only alone (n=95, 18 R / 77 NR), the limited single-cohort sample size produces a wide effect-size CI ($g$ = 0.01, 95% CI: −0.50 to 0.52; TOST $p$ = 0.23). The 22-cohort meta-analytic estimate provides substantially greater precision: pooled $g$ = 0.076 (95% CI: −0.082 to 0.233, SE = 0.081; TOST $p$ = 0.061). The TOST $p$-value narrowly misses the conventional 0.05 threshold for formal equivalence at margin $|g| \leq 0.20$; however, the upper 95% CI bound rules out effects larger than $g$ = 0.23, indicating that any residual association of the signature with response in IO-only settings is, at most, of marginal magnitude. Implementation: `pipeline/11_equivalence_tests.R` (output saved to `output/11_equivalence_tests.rds`).

#### 2.4 Comparison with Known Immune Signatures

Five established immune-activation and immunotherapy-response signatures were tested under identical conditions on both IMvigor210 subgroups (Supplementary Figure S8A; Supplementary Table S6):

- IFN-$\gamma$ signature
- Tumor Inflammation Signature (TIS)
- Gene Expression Profile (GEP)
- T-effector signature
- IPRES resistance signature

In contrast to the chemo-conversion signature, canonical immune signatures either predicted response in both subgroups or neither, consistent with measuring baseline immune status rather than chemotherapy-induced conversion potential. Only the chemo-conversion signature exhibited the interaction pattern (significant in Chemo+IO, non-significant in IO-only).

#### 2.5 Baseline Immune Marker Correlations

Spearman correlations between baseline chemo-conversion signature scores and immune-hot markers were uniformly weak across all validation cohorts (Supplementary Figure S8B; Supplementary Table S12). These results demonstrate that the chemo-conversion signature does not recapitulate immune-hot status, but rather captures a distinct transcriptomic program related to the propensity for chemotherapy-induced TME remodeling.

219 # 3 Supplementary Figures

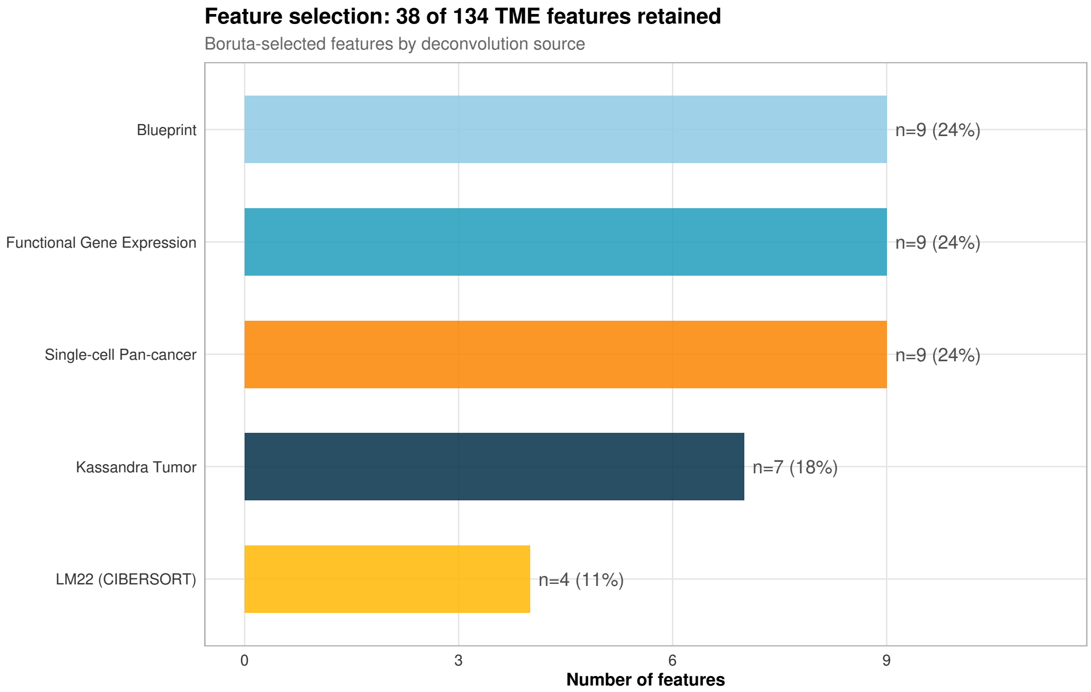

**Figure S1: Boruta-selected feature composition.** Source breakdown of the 38 TME features retained by Boruta feature selection (of 134 initial features), by deconvolution reference: Blueprint (9), functional gene expression (9), single-cell pan-cancer (9), Kassandra tumor (7), and LM22/CIBERSORT (4). Boruta was chosen because it achieved the highest out-of-bag AUC (0.639) among the 10 feature-selection strategies evaluated.

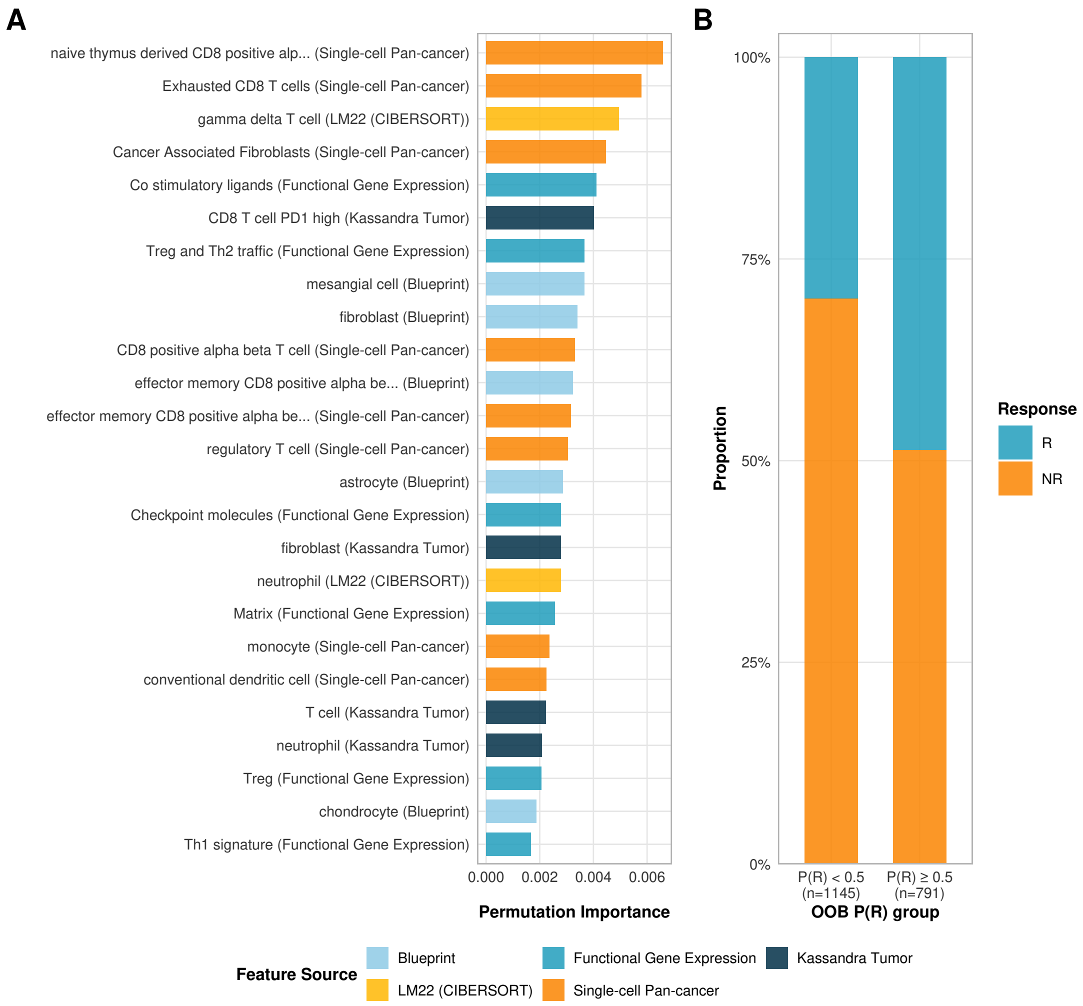

**Figure S2: Random forest model. (A)** The 25 most predictive TME features by permutation importance, colored by deconvolution reference source. **(B)** Out-of-bag calibration: observed responder/non-responder proportions among training samples, split by predicted favorability (P(R) $< 0.5$, n $= 1{,}145$; P(R) $\geq 0.5$, n $= 791$). The model achieved an out-of-bag AUC of 0.639 and a median leave-one-dataset-out AUC of 0.654 across 5 cancer-type folds.

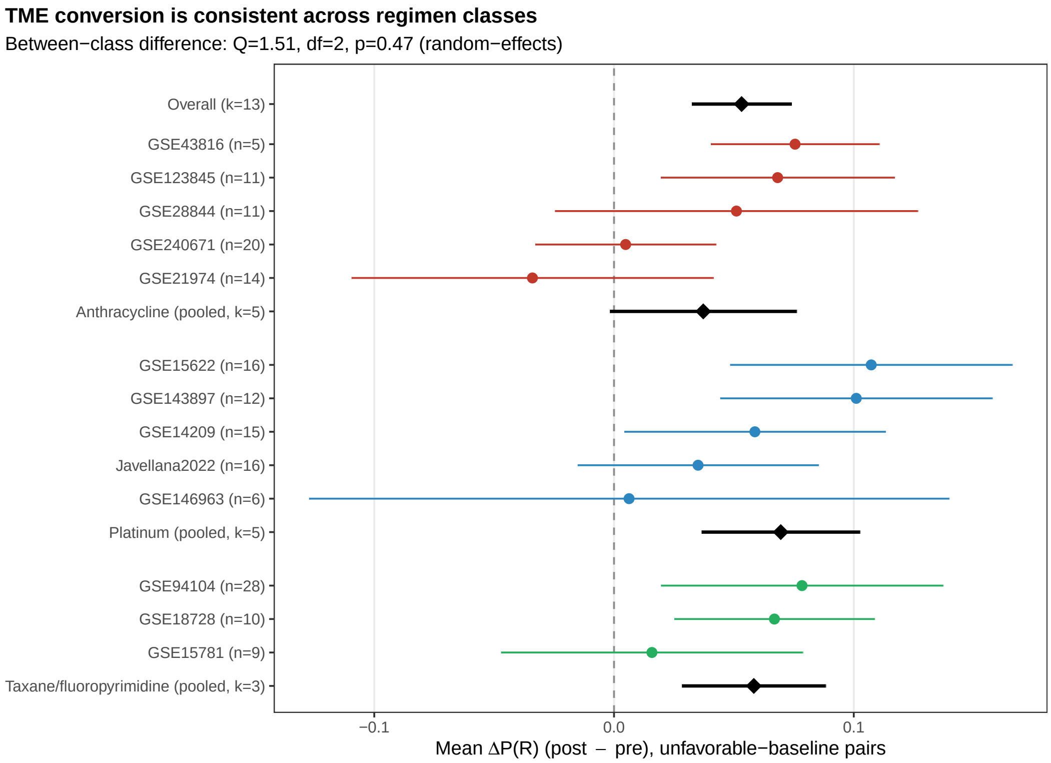

**Figure S3: Chemotherapy-induced TME conversion is consistent across chemotherapy regimen classes.** Forest plot of the per-dataset mean shift in predicted response probability, $\Delta P(R) = P(R)_{post} - P(R)_{pre}$, computed over unfavorable-baseline pairs ($P(R)_{pre} \leq 0.50$) in each of the 13 paired-chemotherapy discovery datasets (points, 95% CI). Because $\Delta P(R)$ depends only on the random-forest TME-favorability axis and not on the signature genes, it is an independent (non-circular) read-out of conversion. Datasets are grouped by an immunogenic-cell-death (ICD)-relevant regimen class assigned from curated drug names: *Anthracycline-based* (doxorubicin/epirubicin ± taxane; high ICD), *Platinum-based* without anthracycline (cisplatin/carboplatin ± taxane; moderate ICD), and *Taxane/fluoropyrimidine* (taxane and/or capecitabine/5-FU; low ICD). Black diamonds are random-effects pooled estimates per class and overall. Conversion is significant overall (pooled $\Delta P(R) = 0.053$, 95% CI: 0.032–0.074, $p < 0.001$) and positive in every regimen class, with no significant difference between classes (moderator test $Q = 1.51$, df $= 2$, $p = 0.47$). Notably the effect is *not* concentrated in the high-ICD anthracycline regimens, indicating that chemotherapy-induced conversion is a general property of cytotoxic chemotherapy rather than an anthracycline-specific effect. Per-dataset values are listed in Supplementary Table S3.

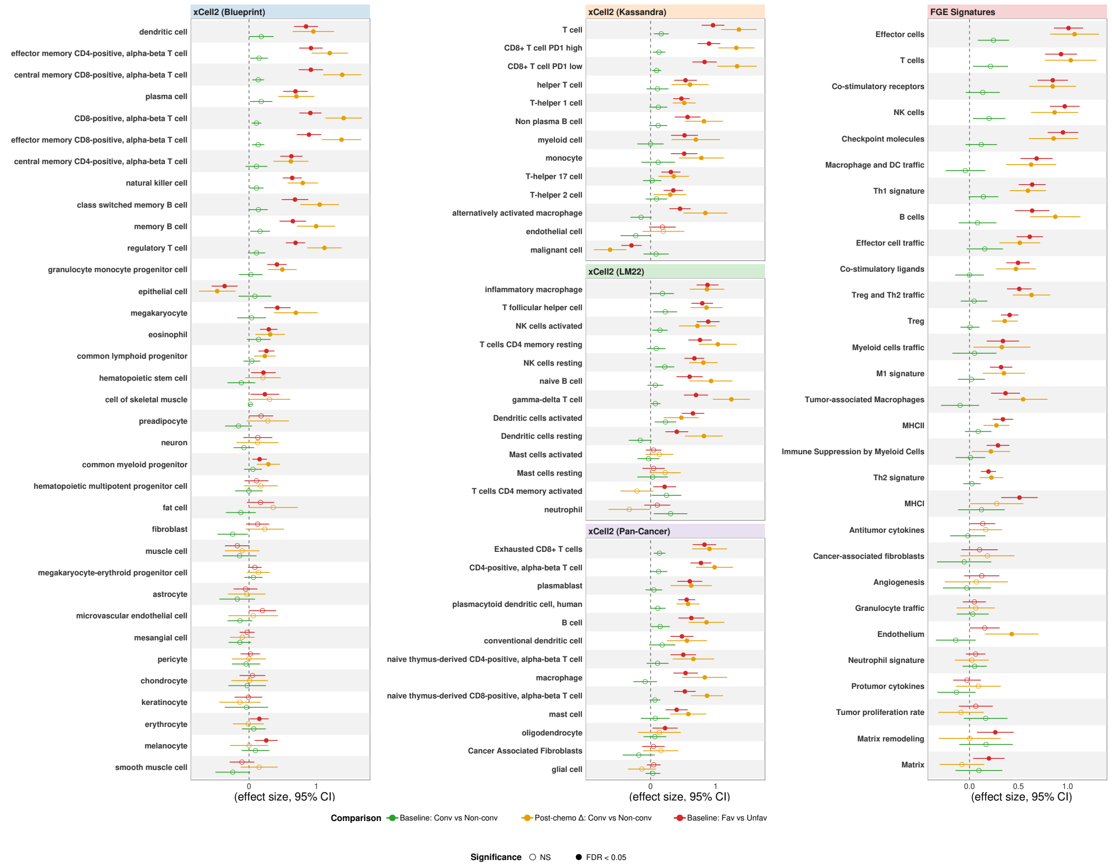

**Figure S4: Full TME remodeling analysis.** Mixed-effect model effect sizes (95% CI) for all 103 TME features across three comparisons: baseline converter vs. non-converter, post-chemotherapy Δ converter vs. non-converter, and baseline favorable vs. unfavorable. Filled circles: FDR $< 0.05$.

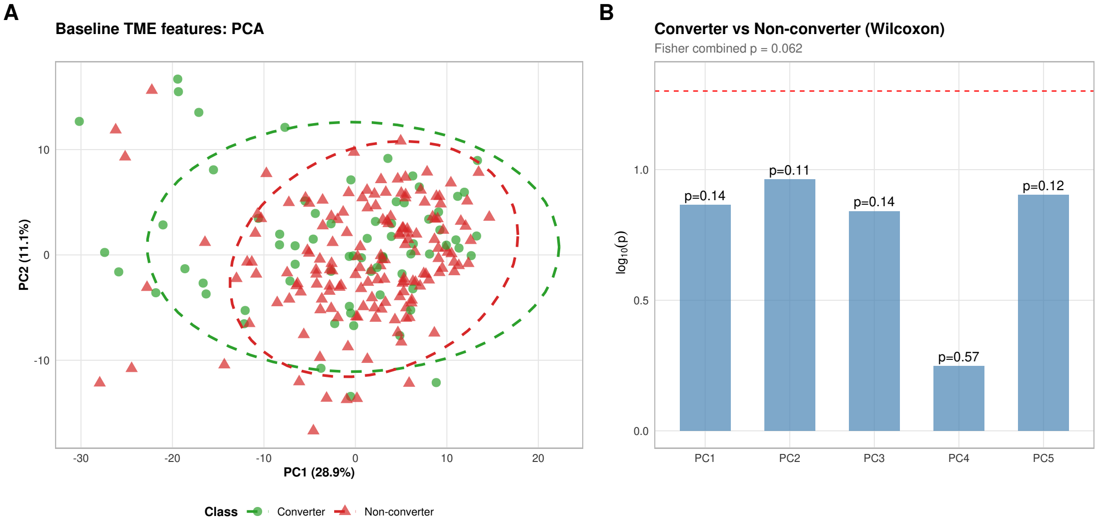

**Figure S5: PCA of baseline TME features.** Principal component analysis on all 103 per-dataset-normalized TME features shows no significant separation between converters and non-converters along any of the top five principal components (all Wilcoxon p > 0.1).

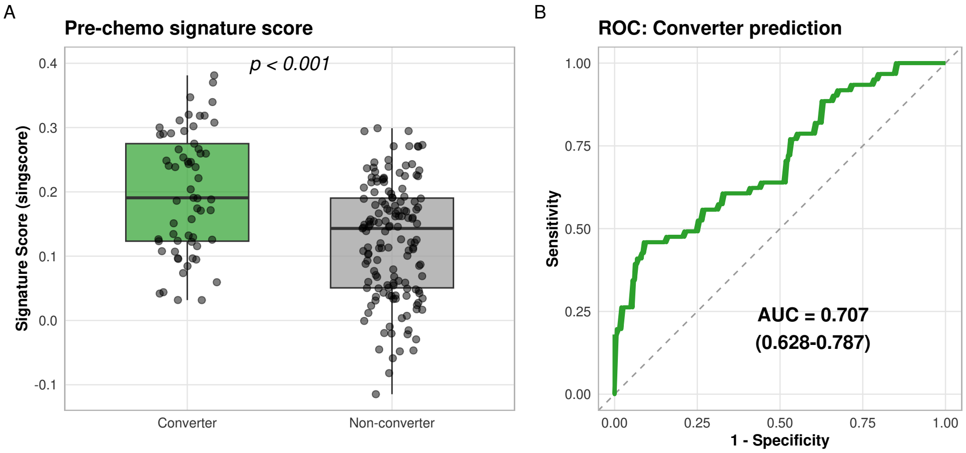

**Figure S6: Baseline signature score separation of converters and non-converters in discovery cohorts. (A)** Distribution of chemo-conversion signature scores at baseline in converters ($n = 61$) vs. non-converters ($n = 148$ classified, of which 147 had baseline expression data and are shown) from the paired chemotherapy discovery cohorts. **(B)** ROC curve for converter classification by baseline signature score (AUC = 0.707, 95% CI: 0.628–0.787). This analysis represents an internal consistency check, as the signature was derived from differential expression between these same groups.

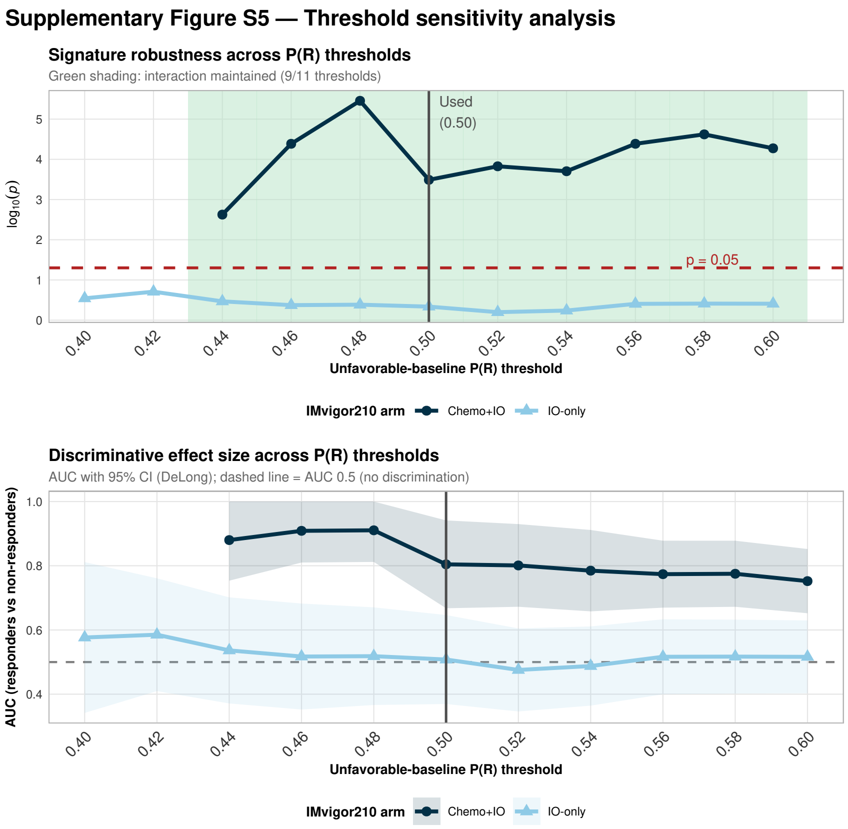

**Figure S7: Threshold sensitivity analysis. (A)** Wilcoxon p-values for signature score association with response across P(R) thresholds (0.40–0.60) in the IMvigor210 Chemo+IO (blue) and IO-only (orange) subgroups. **(B)** Discriminative effect size (AUC with 95% DeLong CI) per arm across the same thresholds; the dashed line marks AUC = 0.5 (no discrimination). The Chemo+IO AUC stays well above 0.5 across thresholds while the IO-only AUC remains near 0.5 throughout.

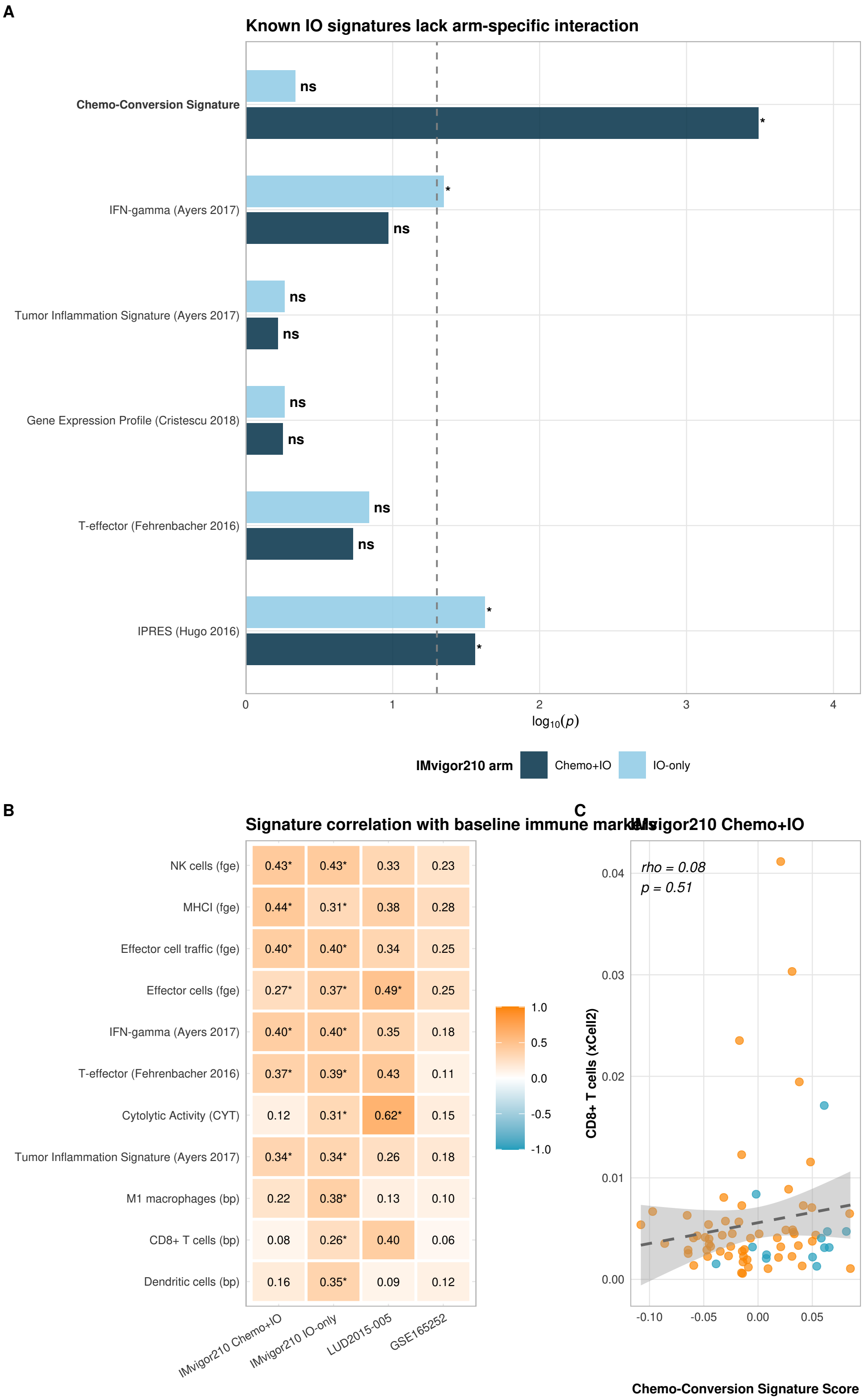

**Figure S8: The chemo-conversion signature is orthogonal to baseline immune status.** **(A)** Comparison of known immunotherapy-response signatures across both IMvigor210 treatment arms. **(B)** Spearman correlation heatmap between baseline chemo-conversion signature scores and immune markers across validation cohorts. **(C)** Representative scatter plots of signature score versus baseline $CD8^{+}$ T cell abundance in both IMvigor210 arms.

### 4 Supplementary Tables

**Table S2: Paired chemotherapy discovery datasets.** 19 datasets, 334 paired patients (209 unfavorable-baseline; 61 converters, 148 non-converters).

| Dataset | Cancer | Platform | Pairs | Drug Class |
|---|---|---|---|---|
| GSE14209 | Gastric | Microarray | 22 | DNA disruption |
| GSE21974 | Breast | Microarray | 25 | DNA disruption |
| GSE28844 | Breast | Microarray | 28 | DNA disrupt + MT disrupt |
| GSE43816 | Breast | Microarray | 7 | DNA disrupt + MT disrupt |
| GSE55410 | Ovarian | Microarray | 9 | DNA disruption |
| GSE15622 | Ovarian | Microarray | 29 | MT disrupt / DNA disrupt |
| GSE146963 | Ovarian | Microarray | 9 | DNA disrupt + MT disrupt |
| GSE18728 | Breast | Microarray | 16 | DNA disruption |
| GSE15781 | Colorectal | Microarray | 9 | Chemotherapy |
| GSE94104 | Colorectal | Microarray | 40 | Chemotherapy |
| GSE32072 | Breast | Microarray | 21 | Chemotherapy |
| GSE102118 | Ovarian | RNA-Seq | 8 | DNMTi + DNA disrupt |
| GSE143897 | Ovarian | RNA-Seq | 18 | DNA disrupt + MT disrupt |
| GSE123845 | Breast | RNA-Seq | 19 | Chemotherapy |
| GSE180280 | Breast | RNA-Seq | 3 | Chemotherapy |
| GSE191127 | Breast | RNA-Seq | 20 | DNA disruption |
| GSE213065 | Sarcoma | RNA-Seq | 4 | DNA disrupt (chemo only) |
| GSE240671 | Breast | RNA-Seq | 27 | Chemotherapy |
| Javellana2022 | Ovarian | RNA-Seq | 20 | Chemotherapy |
| **Total** | | | **334** | |

**Table S3: Chemotherapy-induced conversion by regimen class.** Per-dataset mean $\Delta$P(R) ($P(R)_{post} - P(R)_{pre}$, unfavorable-baseline pairs) for the 13 paired-chemotherapy discovery datasets, by ICD-relevant regimen class, as in Supplementary Figure S3. Available as CSV: `tables/supplementary/stable13_regimen_conversion.csv`.

**Table S4: Chemo-conversion signature gene list.** Complete 50-gene signature (29 up, 21 down) ordered by Robust Rank Aggregation score; all genes pass RRA < 0.01.

| Gene | Direction | RRA score |
|---|---|---|
| *KLK10* | Up | 5.7e-07 |
| *LAGE3* | Up | 8.7e-07 |
| *RUNX3* | Up | 1.2e-06 |
| *CORO1A* | Up | 1.8e-06 |
| *GLRX5* | Up | 2.2e-06 |
| *KLK11* | Up | 2.7e-06 |
| *CLDN4* | Up | 3.7e-06 |
| *PLCE1* | Up | 3.9e-06 |
| *DAG1* | Up | 6.3e-06 |
| *TAOK3* | Up | 7.9e-06 |
| *ADNP* | Up | 9.6e-06 |
| *KRT14* | Up | 1.0e-05 |
| *RRS1* | Up | 1.0e-05 |
| *FEN1* | Up | 1.1e-05 |
| *ZBP1* | Up | 1.1e-05 |
| *UFD1* | Up | 1.2e-05 |
| *UTP18* | Up | 1.2e-05 |
| *CD8A* | Up | 1.3e-05 |
| *STK24* | Up | 1.3e-05 |
| *CCL5* | Up | 1.3e-05 |
| *MRM2* | Up | 1.4e-05 |
| *SNAP29* | Up | 1.6e-05 |
| *FAM53B* | Up | 1.6e-05 |
| *EPOP* | Up | 1.7e-05 |
| *CKB* | Up | 1.9e-05 |
| *CD27* | Up | 2.1e-05 |
| *TNF* | Up | 2.1e-05 |
| *SPAG4* | Up | 2.2e-05 |
| *PTPRC* | Up | 2.2e-05 |
| *ECM1* | Down | 8.1e-07 |
| *LRRC17* | Down | 1.4e-06 |

*Continued on next page*

*(continued from previous page)*

| **Gene** | **Direction** | **RRA score** |
|---|---|---|
| *CNTNAP1* | Down | 1.9e-06 |
| *SORBS3* | Down | 6.8e-06 |
| *SEMA6A* | Down | 9.7e-06 |
| *RGS4* | Down | 1.1e-05 |
| *SLC14A1* | Down | 1.2e-05 |
| *EPHA7* | Down | 1.2e-05 |
| *DUSP4* | Down | 1.3e-05 |
| *COL4A5* | Down | 1.4e-05 |
| *RUSC2* | Down | 1.6e-05 |
| *CYP3A5* | Down | 1.8e-05 |
| *KRTAP5-9* | Down | 1.9e-05 |
| *CAMK1* | Down | 2.2e-05 |
| *CAVIN2* | Down | 3.0e-05 |
| *STK39* | Down | 3.0e-05 |
| *SETBP1* | Down | 3.2e-05 |
| *NOVA1* | Down | 3.7e-05 |
| *ALX4* | Down | 3.8e-05 |
| *LAMC3* | Down | 3.8e-05 |
| *NCSTN* | Down | 3.9e-05 |

**Table S5: IMvigor210 clinical baseline characteristics by treatment trajectory.** Baseline clinical and molecular characteristics of the unfavorable-baseline IMvigor210 interaction population (P(R) $\leq 0.50$), by the three treatment trajectories (Chemo+IO; IO-only naive; IO-only post-chemo biopsy). Only ECOG differed across trajectories (Fisher $p < 0.001$); the chemo-conversion signature score itself did not ($p = 0.561$).

| Characteristic | Level | Chemo+IO (n=72) | IO-only naive (n=38) | IO-only post-chemo (n=57) | *p* |
|---|---|---|---|---|---|
| Response | R | 13 (18.1%) | 11 (28.9%) | 7 (12.3%) | 0.127 |
| | NR | 59 (81.9%) | 27 (71.1%) | 50 (87.7%) | |
| ECOG performance status | 0 | 38 (52.8%) | 10 (26.3%) | 21 (36.8%) | <0.001 |
| | 1 | 34 (47.2%) | 20 (52.6%) | 35 (61.4%) | |
| | 2 | 0 (0.0%) | 8 (21.1%) | 1 (1.8%) | |
| Metastatic disease site | Liver | 19 (29.2%) | 10 (29.4%) | 20 (35.7%) | 0.927 |
| | LN Only | 8 (12.3%) | 5 (14.7%) | 7 (12.5%) | |
| | Visceral | 38 (58.5%) | 19 (55.9%) | 29 (51.8%) | |
| Sex | F | 15 (20.8%) | 10 (26.3%) | 13 (22.8%) | 0.800 |
| | M | 57 (79.2%) | 28 (73.7%) | 44 (77.2%) | |
| Race | ASIAN | 1 (1.4%) | 2 (5.3%) | 1 (1.8%) | 0.596 |
| | BLACK OR AFRICAN AMERICAN | 3 (4.2%) | 2 (5.3%) | 1 (1.8%) | |
| | OTHER | 3 (4.2%) | 0 (0.0%) | 0 (0.0%) | |
| | UNKNOWN | 1 (1.4%) | 0 (0.0%) | 1 (1.8%) | |
| | WHITE | 64 (88.9%) | 34 (89.5%) | 54 (94.7%) | |
| Tobacco use history | CURRENT | 10 (13.9%) | 2 (5.3%) | 8 (14.0%) | 0.647 |
| | NEVER | 22 (30.6%) | 13 (34.2%) | 20 (35.1%) | |
| | PREVIOUS | 40 (55.6%) | 23 (60.5%) | 29 (50.9%) | |
| Immune phenotype | desert | 24 (38.7%) | 11 (36.7%) | 23 (56.1%) | 0.365 |
| | excluded | 32 (51.6%) | 15 (50.0%) | 14 (34.1%) | |
| | inflamed | 6 (9.7%) | 4 (13.3%) | 4 (9.8%) | |
| TCGA subtype | I | 44 (61.1%) | 15 (39.5%) | 23 (40.4%) | 0.048 |
| | II | 11 (15.3%) | 11 (28.9%) | 8 (14.0%) | |
| | III | 8 (11.1%) | 8 (21.1%) | 12 (21.1%) | |
| | IV | 9 (12.5%) | 4 (10.5%) | 14 (24.6%) | |
| Tumor mutation burden /MB | median [IQR] | 7.00 [5.00, 11.00] | 6.50 [4.00, 13.25] | 8.00 [5.00, 12.00] | 0.956 |
| Neoantigen burden /MB | median [IQR] | 0.75 [0.43, 1.45] | 0.75 [0.29, 1.33] | 0.73 [0.43, 1.26] | 0.787 |
| Baseline P(R), model | median [IQR] | 0.44 [0.40, 0.47] | 0.40 [0.37, 0.44] | 0.41 [0.32, 0.45] | 0.003 |
| Chemo-conversion signature score | median [IQR] | −0.01 [−0.03, 0.03] | −0.01 [−0.03, 0.02] | 0.00 [−0.04, 0.05] | 0.561 |
| Overall survival (months) | median [IQR] | 10.48 [5.31, 20.57] | 13.09 [5.39, 17.06] | 7.89 [2.66, 12.85] | 0.005 |

**Table S6: Interaction comparison with known immune signatures.** Wilcoxon $p$-values and AUC for five established immune signatures in both IMvigor210 arms. Available as CSV: `tables/supplementary/stable8_interaction_comparison.csv`.

**Table S7: Cancer-type-balanced LODO folds.** The five held-out folds used for cancer-type-balanced leave-one-dataset-out (LODO) hyperparameter selection. Each fold holds out the largest cohort of one cancer type while the model is trained on the remaining 26 cohorts (1,936 samples total); the selected configuration achieved a median held-out AUC of 0.654.

| **Held-out cancer-type fold** | **Held-out samples ($n$)** |
|---|---|
| Melanoma | 124 |
| RCC | 380 |
| NSCLC | 129 |
| HNSCC | 102 |
| Gastric | 45 |

**Table S8: Random forest model performance metrics.** Final random-forest classifier evaluated out-of-bag (OOB) at the Youden-optimal probability threshold (0.470); the cancer-type-balanced LODO median AUC was 0.654 (Table S7).

| **Metric** | **Value** |
|---|---|
| OOB AUC | 0.639 |
| Optimal threshold (Youden $J$) | 0.470 |
| Sensitivity | 66.8% |
| Specificity | 56.0% |
| Accuracy | 60.1% |
| Positive predictive value | 47.8% |
| Negative predictive value | 73.7% |
| $N$ samples (OOB) | 1,936 |

**Table S9: Signature–marker gene overlap test.** Overlap of the 50 signature genes with the xCell2 reference panels (12,612 genes) and 29 FGE pathway sets (243 genes). Three genes (CD8A, CD27, TNF) overlap the FGE sets; 42 overlap the broader panels (Fisher OR = 13.0, $p < 0.001$), reflecting compendium breadth, not redundancy.

**Table S10: Validation results (AUC and two-sided $p$-values).** Two-sided Wilcoxon $p$-values with DeLong 95% CIs for AUC across IMvigor210 strata and the five validation cohorts.

| Cohort | n (R/NR) | AUC (95% CI) | $p_{\text{two-sided}}$ |
|---|---|---|---|
| IMvigor210 Chemo+IO Unfav | 13/59 | 0.804 (0.668–0.941) | 0.00065 |
| IMvigor210 IO-only Unfav | 18/77 | 0.508 (0.369–0.646) | 0.92 |
| IMvigor210 Chemo+IO Fav | 18/42 | 0.630 (0.485–0.774) | 0.13 |
| IMvigor210 IO-only Fav | 17/48 | 0.494 (0.316–0.672) | 0.95 |
| LUD2015-005 Unfav | 12/6 | 0.847 (0.654–1.000) | 0.018 |
| GSE165252 Unfav | 10/18 | 0.733 (0.545–0.922) | 0.045 |
| GSE207422 Unfav | 3/10 | 0.633 (0.086–1.000) | 0.57 |
| GSE241876 Unfav | 6/3 | 0.722 (0.342–1.000) | 0.38 |
| TONIC Chemo+IO Unfav | 4/5 | 0.650 (0.206–1.000) | 0.56 |

**Table S11: ICD pillar gene set assignments.** 13 immunogenic-cell-death gene sets organized by Kroemer et al.'s four-pillar framework. Available as XLSX: `tables/supplementary/icd_gene_sets.xlsx`.

**Table S12: Immune marker correlation table.** Spearman $\rho$ and $p$-values between baseline signature scores and immune markers across validation cohorts. Available as CSV: `tables/supplementary/stable9_immune_marker_correlations.csv`.

**Table S13: IMvigor210 arm TME feature comparison.** Comparison of the 38 Boruta-selected TME features between Chemo+IO (n=72) and IO-only (n=95) unfavorable-baseline subgroups; none differed (0/38 at FDR $< 0.05$). Available as CSV: `tables/stable10_imvigor210_arm_tme.csv`.

### 5 Reporting Guideline Checklists

This study both develops a machine-learning prediction model (a random-forest classifier of tumor-microenvironment favorability) and derives and evaluates a predictive transcriptomic biomarker (the 50-gene chemo-conversion signature). It is therefore reported in accordance with the REMARK guideline for tumor-marker prognostic studies (McShane et al., 2005) and the TRIPOD+AI guideline for prediction-model studies using machine learning (Collins et al., 2024). Checklist item text below is abbreviated; the "Reported in" column points to the relevant sections, figures, and tables of the main text and this Supplement. Item wording should be confirmed against the official published checklists before submission.

**Table S14: REMARK checklist** (REporting recommendations for tumor MARKer prognostic studies; McShane et al., 2005).

| # | Checklist item (abbreviated) | Reported in |
|---|---|---|
| 1 | Marker examined, study objectives, pre-specified hypotheses | Abstract; Introduction (final paragraph) |
| 2 | Patient characteristics, source, inclusion/exclusion criteria | Methods (Data Collection); Supp. Methods (Dataset Descriptions); Tables S1–S2 |
| 3 | Treatments received and how chosen | Methods (Data Collection); Supp. Methods; Tables S1–S2 |
| 4 | Biological material; methods of preservation/storage | Methods (Data Collection: RNA-Seq TPM, Affymetrix RMA); Supp. Methods |
| 5 | Assay method and protocol; quality control; blinding | Methods (TME Feature Quantification; Signature Scoring); Supp. Methods (singscore, RRA). Scoring is a pre-specified computational pipeline applied uniformly |

*Continued on next page*

*(REMARK checklist, continued)*

| # | Checklist item (abbreviated) | Reported in |
|---|---|---|
| 6 | Case selection (prospective/retrospective); time period; follow-up | Retrospective secondary analysis of public cohorts (Methods). Response is cross-sectional; overall survival is exploratory (Discussion); median follow-up not applicable to the primary endpoint |
| 7 | Clinical endpoints precisely defined | Methods (Data Collection): responder (CR/PR) vs non-responder (SD/PD) per RECIST v1.1; overall survival (IMvigor210) |
| 8 | All candidate variables initially examined | Methods (144 TME features → 134 after filtering → 38 Boruta-selected) |
| 9 | Sample-size rationale / power | Retrospective design; no a priori sample-size calculation. Post-hoc power for small validation cohorts and imprecision discussed (Results; Discussion) |
| 10 | Statistical methods, variable selection, assumptions, missing data | Methods (Statistical Analysis; RF/Boruta; RRA; Wilcoxon; REML meta-analysis; TOST); per-dataset normalization and replicate handling (Methods; Supp. Methods) |
| 11 | Handling of marker values; cutpoints | Methods (continuous singscore; unfavorable baseline $P(R) \leq 0.50$; median split for the odds-ratio forest) |

*Continued on next page*

*(REMARK checklist, continued)*

| # | Checklist item (abbreviated) | Reported in |
|---|---|---|
| 12 | Flow of patients; numbers and events at each stage | Results; Figures 1 and 2A (209 unfavorable-baseline; 61 converters / 148 non-converters; IMvigor210 72 Chemo+IO / 95 IO-only); Tables S2, S5 |
| 13 | Distributions of demographics, prognostic variables, marker; missing values | Supp. Table S5 (IMvigor210 clinical baseline by trajectory); Results |
| 14 | Relation of marker to standard prognostic variables | Results (signature score vs ECOG, metastatic site, mutational/neoantigen burden; balanced, $p$=0.56); Supp. Table S5 |
| 15 | Univariable marker–outcome effect with CI (KM if time-to-event) | Results (per-cohort AUC, one-sided Wilcoxon, Hedges' $g$); Figure 3; KM survival Figure 3B; Supp. Table S10 |
| 16 | Key multivariable analyses; effects with CI | Results (logistic-regression signature×arm interaction, OR with 95% CI; ECOG-adjusted) |
| 17 | Estimated effects with CI regardless of significance | Results; Supp. Table S10 (all strata and cohorts: AUC, two-sided $p$, CI) |
| 18 | Interpretation in context of hypotheses and other studies | Discussion |
| 19 | Discussion of limitations (bias, imprecision, generalizability) | Discussion (Limitations) |
| 20 | Implications for future research and clinical value | Discussion (clinical promise; future work) |

**Table S15: TRIPOD+AI checklist** (Transparent Reporting of a multivariable prediction model for Individual Prognosis Or Diagnosis — artificial-intelligence extension; Collins et al., 2024).

| # | Checklist item (abbreviated) | Reported in |
|---|---|---|
| 1 | Title identifies the study as developing/evaluating a multivariable (ML) prediction model, with target population and outcome | Title |
| 2 | Structured abstract | Abstract |
| 3 | Background and rationale; existing models/biomarkers and their limitations | Introduction |
| 4 | Objectives (development and external validation) | Introduction (final paragraph) |
| 5 | Source of data and study design | Methods (Data Collection); Supp. Methods. Retrospective multi-cohort analysis of public data |
| 6 | Participants: eligibility, settings | Methods; Supp. Methods (Dataset Descriptions); Tables S1–S2 |
| 7 | Treatments received (relevant to outcome) | Methods; Supp. Methods; Tables S1–S2 |
| 8 | Outcome: definition, how and when assessed; blinding | Methods (responder/non-responder per RECIST v1.1; overall survival). Outcome assessment independent of the computational predictor |
| 9 | Predictors: definition, measurement; preprocessing | Methods (TME Feature Quantification: xCell 2.0 + FGE, 144 features) |
| 10 | Sample size | 1,936 training; 334 paired; per-cohort $n$ (Methods; Tables S1–S2). No a priori calculation (retrospective) |
| 11 | Missing data handling | Methods (replicate collapse; per-dataset normalization; near-zero-MAD filter); Supp. Methods |

*Continued on next page*

*(TRIPOD+AI checklist, continued)*

| # | Checklist item (abbreviated) | Reported in |
|---|---|---|
| 12 | Data preprocessing and predictor preparation | Methods (Batch Effect Correction: per-dataset median-MAD normalization; Boruta feature selection 144→134→38); Supp. Methods |
| 13 | Model type/algorithm and how built (hyperparameter tuning) | Methods (RF, ranger; cancer-type-balanced LODO hyperparameter search) |
| 14 | How the model produces predictions; thresholds | Methods (P(R) probability; singscore signature score; unfavorable baseline P(R) $\leq$ 0.50) |
| 15 | Performance measures | Methods (AUC [pROC]; one-sided Wilcoxon; Hedges' $g$; REML meta-analysis; odds ratios) |
| 16 | Model evaluation: internal and external validation procedures | Methods (out-of-bag + cancer-type-balanced LODO internal; six external validation cohorts; 22 IO-only specificity cohorts) |
| 17 | Class-imbalance handling | Methods (inverse-frequency class weights) |
| 18 | Fairness / subgroup performance | Cross-cancer-type generalization assessed via LODO; demographic-subgroup fairness not formally evaluated (Discussion, Limitations) |
| 19 | Study registration / protocol | Not registered; retrospective secondary analysis of publicly available data |
| 20 | Funding and role of funders | Acknowledgments (Israel Science Foundation 1543/21; funder had no role) |

*Continued on next page*

*(TRIPOD+AI checklist, continued)*

| # | **Checklist item (abbreviated)** | **Reported in** |
|---|---|---|
| 21 | Conflicts of interest | Disclosure of Potential Conflicts of Interest (none) |
| 22 | Data availability | Data Availability (GEO, Synapse, IMvigor210CoreBiologies; TONIC under EGA controlled access) |
| 23 | Code availability | Data Availability (GitHub; Zenodo upon publication) |
| 24 | Participant flow and characteristics; predictor distributions | Results; Figures 1, 2A, 3; Tables S2, S5 |
| 25 | Model specification and performance (development + validation) | Results (50-gene signature, Figures 2D–E; RF OOB AUC 0.639, LODO 0.654; validation AUCs, Figure 3); Tables S4, S10 |
| 26 | Interpretation, comparison with other studies, limitations | Discussion (including Limitations) |
| 27 | Usability, clinical implications, and future work | Discussion (clinical promise; future work) |
